# Dual profiling of DNA modifications with enhancer features during the exit of naive pluripotency

**DOI:** 10.64898/2026.06.12.731885

**Authors:** Marlet Morales-Franco, Baptiste Demaret, Pol Arnau Romero, Hortense Bouvier, Julien Richard Albert, Aaron B. Bogutz, Angélique David, Pierre-Antoine Defossez, Anaïs Flore Bardet, Priscillia Lhoumaud, Maxim V.C. Greenberg

**Affiliations:** Université Paris Cité, CNRS, Institut Jacques Monod, Paris, France; Department of Medical Genetics, Life Sciences Institute, University of British Columbia, Vancouver, British Columbia, Canada; Université Paris Cité, CNRS, Epigenetics and Cell Fate, Paris, France; Institut de Génétique et de Biologie Moléculaire et Cellulaire (IGBMC), Université de Strasbourg, CNRS, INSERM, Université de Strasbourg, Illkirch, France

## Abstract

Cis-regulatory elements, such as enhancers, play an essential role in coordinating gene expression programs during cellular transitions. As such, substantial efforts have been made to characterize enhancer elements, e.g., chromatin accessibility, transcription factor (TF) binding sites, histone post-translational modifications (PTMs), and 5-cytosine DNA methylation and hydroxymethylation (5mC and 5hmC). Elevated 5mC levels are typically correlated with the inactive enhancer state. However, whether 5mC precludes TF binding or is simply a downstream consequence of enhancer decommissioning is difficult to determine. Bulk genomics assays from cell populations fail to fully capture cellular heterogeneity, and single-cell assays often suffer from limited read counts per cell. Here, we leveraged tagmentation-based technologies to assess chromatin features with 5mC simultaneously: Methyl-ATAC and Methyl-CUT&Tag. In addition, we modified the technique to interrogate 5hmC dynamics (hM-ATAC). We employed these techniques during the exit of naive pluripotency in mouse embryonic stem cells (mESCs), which recapitulates the embryonic DNA methylation establishment program. This system withstands the complete absence of DNA methylation and demethylation machinery, allowing us to further dissect the temporal contributions of 5mC and 5hmC. Given the affordability of these techniques, we were able to obtain robust, dynamic single-molecule 5mC/5hmC information during this cellular transition for chromatin accessibility and histone marks in wild-type and mutant conditions. The sum of these data allowed for unprecedented insight into the role that 5mC turnover plays at enhancers during a key window of mammalian embryonic development.

## Introduction

Enhancer elements are short, non-coding DNA sequences located distally from their target genes, where they regulate gene expression in a spatiotemporal manner. Enhancers can be classified based on their chromatin accessibility, transcription factor occupancy, and specific histone modifications. For example, active enhancers are marked by both histone H3 lysine 4 monomethylation (H3K4me1) and lysine 27 acetylation (H3K27ac), while primed or poised enhancers are characterized by H3K4me1^1,2^. The presence of H3K27ac thus serves as a key distinguishing feature between active and primed enhancers. The regulation of gene expression by enhancers largely depends on three main processes: the binding of transcription factors, the recruitment of RNA Polymerase II—which often leads to the production of enhancer RNAs (eRNAs)^3^—and the physical, long-range interaction between the enhancer and its target gene promoter^4^. Notably, many TFs are sensitive to CpG methylation, which can influence their binding affinity^5^. 5-methylcytosine (5mC) is a conserved epigenetic mark in eukaryotes. In mammals, DNA methyltransferases (DNMTs) primarily establish 5mC at CpG dinucleotides—unless otherwise stated, 5mC will refer to methylated CpGs. The removal of 5mC can occur passively through dilution during DNA replication, or actively via the Ten-eleven Translocation (TET) enzymes, which oxidize 5mC to 5-hydroxymethylcytosine (5hmC) and further intermediates, ultimately resulting in demethylation^6^.

In general, 5mC is associated with transcriptional repression. This is particularly evident at CpG-rich regions, known as CpG islands (CGIs). CGIs overlap with approximately two-thirds of mammalian promoters^7^. The vast majority of CGI promoters are devoid of 5mC; however, methylated promoters correlate with robust silencing^8^. Enhancers, on the other hand, are usually not CpG-rich and are primarily located in intergenic and intragenic regions, where CpGs tend to be methylated^9^.

With some key exceptions, CGI promoter hypomethylation remains stable during development^10^, whereas enhancers often exhibit spatiotemporal changes in 5mC levels that correlate with dynamic gene expression patterns^11^. Globally, there is an inverse relationship between enhancer methylation and gene expression, raising important questions about the functional significance of 5mC turnover at enhancers^12^. DNA methylation at enhancers could directly repress transcription or, alternatively, methylation changes might reflect the loss of TF binding^12,13^. Intriguingly, there is evidence that methylated enhancers can still support transcriptional activation, as seen in "bivalent" enhancers where 5mC coexists with the active histone mark H3K27ac^14^.

Measuring the coexistence of different chromatin features and DNA methylation at the single-molecule level presents a significant technical challenge. Current techniques that simultaneously analyze DNA methylation alongside TF binding or chromatin accessibility^15–23^ often face limitations. These include the analysis of naked DNA rather than native chromatin, the need for large amounts of input material, high sequencing costs, and/or limited resolution at the single-molecule level.

Here, we have implemented the EpiMethylTag technique, which enables the simultaneous profiling of chromatin features (e.g., histone modifications, chromatin accessibility) and DNA methylation^24,25^. Its key advantages include single-molecule resolution, low input requirements, preservation of native chromatin context, cost-efficiency, and scalability. Methyl-Assay for Transposase-Accessible Chromatin (M-ATAC) assesses the correlation between DNA methylation and chromatin accessibility, revealing the extent to which these features coincide^24,26,27^. Additionally, Methyl-Cleavage Under Targets and Tagmentation (M-CUT&Tag)^28^ provides an alternative to traditional ChIP-bisulfite sequencing with many of the same benefits as M-ATAC, enabling the simultaneous analysis of histone modifications and DNA methylation^17,18,24,29^.

Although the powerful EpiMethylTag technique opens new avenues for a better understanding of enhancer methylation regulation during early development, the use of bisulfite conversion does not discriminate between 5mC and 5hmC levels. As 5mC is considerably more abundant in the genome compared to 5hmC, the signal observed is often assumed to correspond to 5mC. However, at enhancers where 5mC turnover is high^22,30^, it is important to distinguish between 5mC and 5hmC signals to determine whether chromatin accessibility and histone modifications coexist with 5mC and/or 5hmC. To date, several techniques allow for the detection of 5hmC levels^31–34^. By adapting the EpiMethylTag strategy, we developed hM-ATAC, a new method that detects 5hmC levels simultaneously with chromatin accessibility from the same molecule. This innovation enables a more precise understanding of how 5hmC, 5mC, and chromatin states coexist and interact during epigenetic remodeling.

During early eutherian mammal embryogenesis, the genome undergoes a wave of demethylation, erasing gametic 5mC patterns up to the blastocyst stage (∼20% 5mC). Following implantation, the *de novo* DNA methyltransferases DNMT3A and 3B remethylate the genome, plateauing at ∼75% 5mC in epiblast stem cells^35^. Proper establishment of 5mC patterns is essential for embryonic development, as exemplified by the embryonic lethality observed in *Dnmt* and *Tet* mutants^36–38^.

These early DNA methylome dynamics can be recapitulated in cell culture. Mouse embryonic stem cells (mESCs) derived from the inner cell mass (ICM) of the blastocyst can be cultured in conditions that maintain globally low 5mC levels. Naive mESCs can thereafter be differentiated into epiblast-like cells (EpiLCs), acquiring the globally high 5mC levels seen *in vivo*. The mESC-to-EpiLC differentiation system has the particular advantage of being resilient to mutations in the DNA methylation and demethylation machinery, thus allowing to dissect the role of 5mC/5hmC in the chromatin landscape and enhancer activity during the early stages of embryonic development.

In this resource, we generated M-ATAC, hM-ATAC, and H3K27ac/H3K4me1 M-CUT&Tag data to characterize dynamic, genome-wide DNA methylation at enhancers during the transition from naive to primed pluripotency. We integrated this data with Whole-Genome Bisulfite Sequencing (WGBS) to draw the distinction between the 5mC associated with a specific chromatin state at a given locus versus 5mC at the same locus in the entire population of cells. Using M-ATAC, we highlighted high chromatin accessibility dynamics during mESC-to-EpiLC differentiation, and provided evidence of coexistence between elevated 5mC and chromatin accessibility. M-ATAC experiments in *Dnmt* and *Tet* KO cells showed that the closing of chromatin at dynamic enhancer regions often required DNA methylation and the binding of specific TFs, but that active demethylation by TET enzymes was not globally required for chromatin opening. In addition, using M-CUT&Tag, we observed that H3K4me1 and 5mC are hardly mutually exclusive, and identified a subset of enhancers that simultaneously harbored active chromatin features (ATAC+, H3K4me1+, and H3K27ac+) and elevated 5mC levels (>50%). Using hM-ATAC, we furthermore discovered that active methylated enhancers displayed higher 5hmC levels than other active and primed enhancers, suggesting that a large proportion of the active methylated class are on a trajectory for lower methylation levels. Finally, motif analysis indicated that both methyl-sensitive and methyl-specific DNA binding factors are implicated in proper enhancer regulation.

The rich datasets generated here provide new insights into the epigenetic regulation of enhancers, revealing how DNA methylation and hydroxymethylation contribute to enhancer activity and gene regulation during early development. This approach overcomes the limitations of previous methods and opens new avenues for understanding the functional significance of enhancer methylation dynamics.

## Results

### M-ATAC reveals DNA methylation dynamics at enhancers

To determine the extent to which 5mC coexists with regulatory elements during the embryonic DNA methylation program, we performed M-ATAC^24,26,27^ in a cellular differentiation system that recapitulates the early embryonic blastocyst-to-epiblast transition (Figure 1A,B). Briefly, native chromatin is first tagmented using Tn5, which inserts C-methylated adapters preferentially into open chromatin regions. The tagmented DNA is then bisulfite-converted to distinguish methylated from unmethylated cytosines, followed by PCR amplification (C-methylated adapter sequences being kept intact during the bisulfite conversion) and sequencing. This approach allows for the direct assessment of chromatin accessibility and DNA methylation patterns. Because both are detected from the same molecule, this cost-effective method mitigates the confounding effects of correlating distinct datasets.

**Figure 1.**
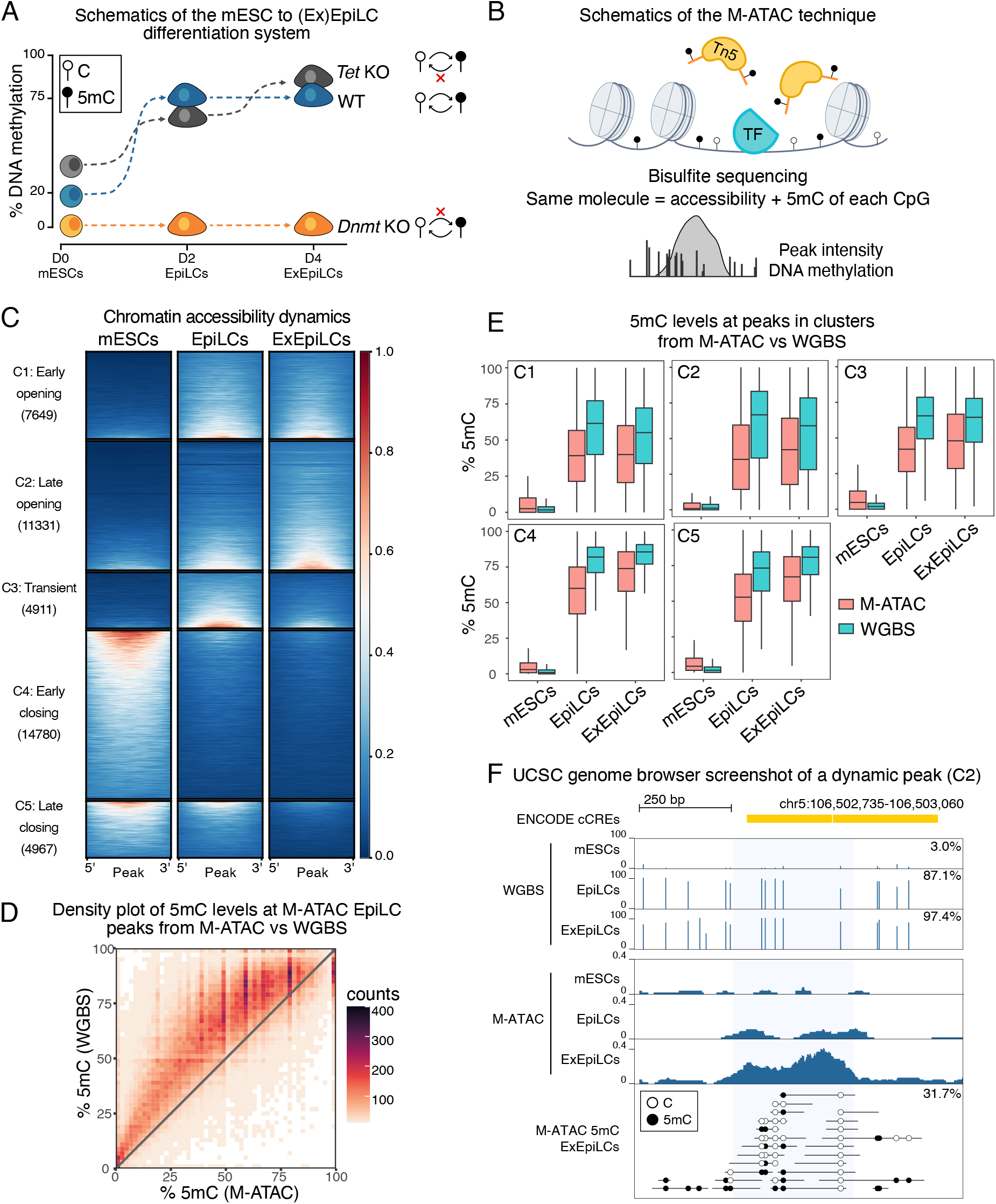
M-ATAC reveals DNA methylation dynamics at enhancers. **A.** Schematics of the 5mC dynamics (Y axis) using a differentiation system from mouse Embryonic Stem Cells (mESCs) to Day 2 Epiblast Like Cells (EpiLCs) to Day 4 Extended EpiLCs (ExEpiLCs). WT cells are in blue, *Dnmt1/3a/3b* triple KO cells (*Dnmt* KO, devoid of DNA methylation) are in orange, and *Tet1/2* double KO cells (*Tet* KO, devoid of active DNA demethylation) are in grey. White lollipops represent unmethylated cytosines and black lollipops represent methylated cytosines (5mC). **B.** Schematic overview of the M-ATAC technique. Tn5 loaded with methylated adaptors "tagment" the DNA at accessible regions. After bisulfite conversion, the adaptors remain Illumina-compatible for high-throughput sequencing. Chromatin accessibility can be assessed simultaneously with the underlying 5mC state. TF: transcription factor. **C.** Heatmaps showing the M-ATAC signal within dynamic non-promoter peaks in WT cells during mESC to EpiLC to ExEpiLC differentiation, divided into 5 clusters: early opening (C1: 7649 peaks), late opening (C2: 11331 peaks), transient peaks (C3: 4911 peaks), early closing (C4: 14780 peaks), and late closing (C5: 4967 peaks). **D.** Density plot of average 5mC from M-ATAC non-promoter peaks compared with WGBS for EpiLCs (91544 peaks, Pearson correlation = 0.78). **E.** Boxplots of 5mC percentages from M-ATAC (red) and WGBS (blue) in mESCs, EpiLCs, and ExEpiLCs, for the 5 clusters presented in panel C. **F.** Representative UCSC genome browser screenshot of M-ATAC and WGBS at a putative enhancer region (ENCODE candidate Cis-Regulatory Elements, cCREs). 5mC level from WGBS is shown on top (mESCs, EpiLCs, and ExEpiLCs, top to bottom), M-ATAC signal is shown in the middle (mESCs, EpiLCs, and ExEpiLCs, top to bottom), and 5mC signal of individual reads from ExEpiLCs is shown at the bottom. White lollipops represent unmethylated cytosines and black lollipops represent methylated cytosines. Blue shading indicates the putative enhancer region defined by our analysis. Methylation percentages of the highlighted region are indicated in the boxes.

We assessed DNA methylation turnover at enhancers using M-ATAC in mESCs cultured with double inhibition of extracellular signal-regulated kinase (ERK) and glycogen synthase kinase (GSK) 3β supplemented with vitamin C (2i+vitC)^39^ and in EpiLCs^40^. mESCs cultured in this media represent the “naive” pluripotent state, and exhibit extreme global DNA hypomethylation^41^. EpiLCs are formed after 2 days of differentiation with FGF2 and Activin A, and recapitulate a transitional state called “formative” pluripotency, during which the global establishment of 5mC occurs. At 4 days of differentiation, Extended EpiLCs (ExEpiLCs) are a closer approximation of the “primed” pluripotent state.

M-ATAC was performed in duplicates in mESCs (day 0), EpiLCs (day 2), and ExEpiLCs (day 4). Following genomic annotation analysis (Figure S1A), peaks that were found at promoter regions (within 3kb of a transcription start site) were removed from further analysis to focus on candidate enhancer regions, with the caveat that some promoters can also behave as enhancers^42^. K-means clustering analysis on regions with dynamic accessibility identified five major clusters: early opening (C1; 7649 peaks), late opening (C2; 11331 peaks), transient (C3; 4911 peaks), early closing (C4; 14780 peaks), and late closing (C5; 4967 peaks) (Figure 1C). The rest of the peaks (84186) were static across time points (Figure S1B). The level of 5mC obtained from M-ATAC peaks was compared to that measured by WGBS. Overall and for each individual cluster, the level of DNA methylation was higher in WGBS than in M-ATAC (Figure 1D-F and Figure S1C-F). The closing regions (clusters 4 and 5) harbored higher 5mC levels than opening regions (clusters 1 and 2) in (Ex)EpiLCs (Figure 1E and Figure S1D), consistent with the known anticorrelation between chromatin accessibility and 5mC. However, we observed intermediate 5mC levels from M-ATAC for every cluster, suggesting that chromatin accessibility could coexist with 5mC globally. In sum, the discordance between WGBS data obtained from a mixed cell population compared with M-ATAC (which exclusively detects 5mC from cells with accessible chromatin) (Figure 1F, Figure S1F), highlights how correlative analyses between WGBS and ATAC datasets can be potentially misleading.

### Mutants reveal 5mC-sensitive enhancers

To assess the role of 5mC in shaping the enhancer landscape, we used mESCs that lack the three embryonic DNMT proteins (*Dnmt1,3a*,*3b* triple knockout; *Dnmt* KO) (complete absence of 5mC)^43^ and mESCs mutant for the TET1 and TET2 proteins (*Tet1*,*2* double knockout; *Tet* KO). The third TET protein, TET3, exhibits low expression in this system^44^. Importantly, *Dnmt* KO and *Tet* KO cells are able to differentiate to EpiLCs^43–46^. In order to determine how DNA methylation or demethylation processes impact enhancer dynamics, we performed M-ATAC in *Dnmt* KO and *Tet* KO mESCs, EpiLCs, and ExEpiLCs (Figure 2A, Figure S2A). Differential accessibility analysis between WT and *Dnmt* KO at peaks within clusters showed broad misregulation of chromatin accessibility. In mESCs and EpiLCs, there were 2010 and 1157 more accessible and 2189 and 1208 less accessible regions in *Dnmt* KO compared to WT, respectively (Figure S2B). Differential peaks in mESCs could be related to the low—but quantifiable—5mC levels in WT, as clusters 1 and 2 (early and late opening, respectively) exhibited precocious accessibility in *Dnmt* KO.

**Figure 2.**
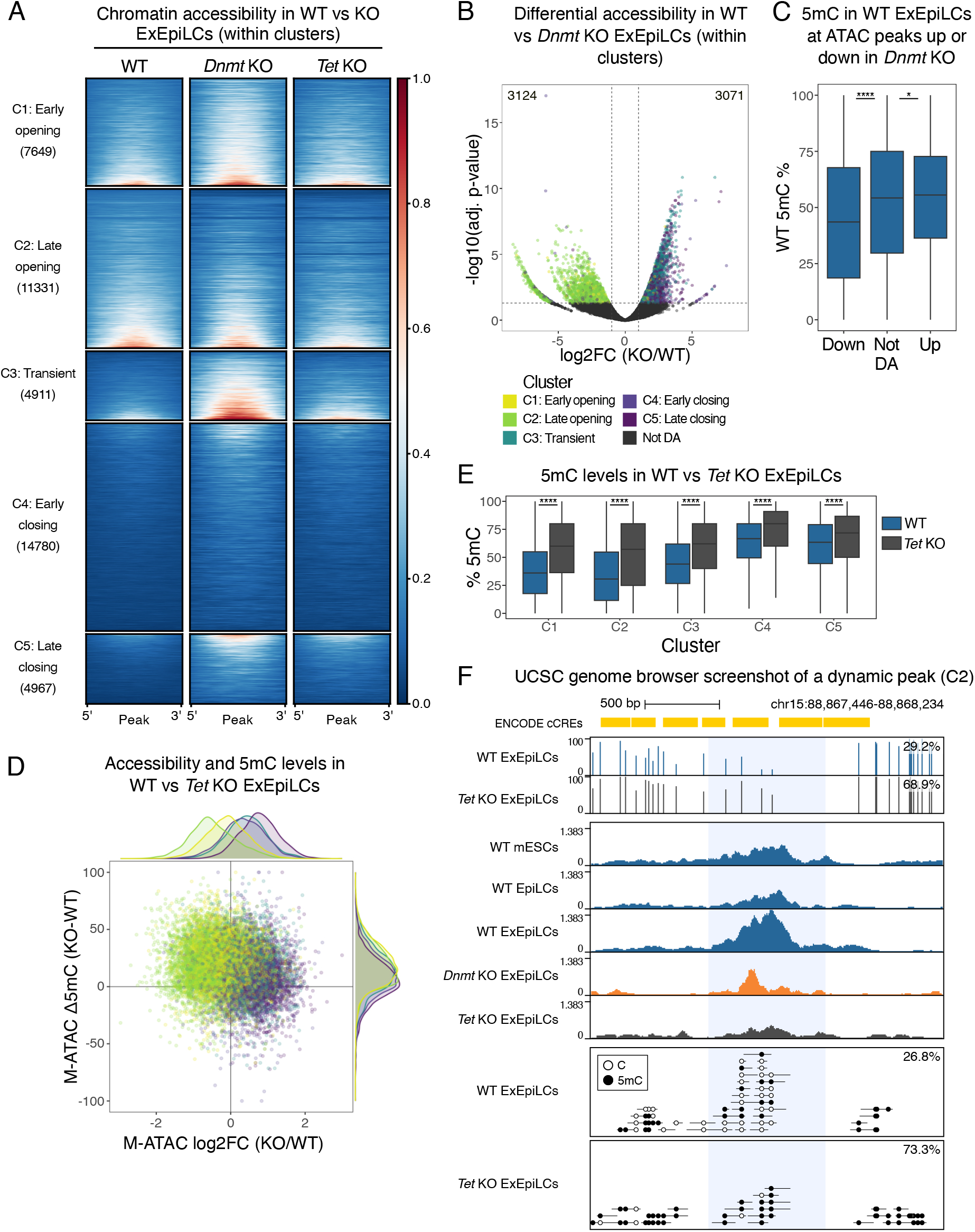
Characterization of *Dnmt* KO and *Tet* KO M-ATAC. **A.** Heatmaps showing the M-ATAC signal in WT, *Dnmt* KO, and *Tet* KO ExEpiLCs, within the 5 clusters defined in Figure 1C. **B.** Volcano plot of differential chromatin accessibility between WT and *Dnmt* KO ExEpiLCs at peaks within the 5 clusters (log2-fold change > 1 or < -1; adjusted p-value < 0.05). Peaks are colored based on their cluster affiliation. The number of differentially accessible peaks is indicated. **C.** Boxplots of 5mC levels in WT ExEpiLCs, comparing the peaks that are less accessible (down), more accessible (up), or not differentially accessible (not DA) in *Dnmt* KO compared to WT ExEpiLCs. **D.** Scatterplot representing the log2 fold change of chromatin accessibility in *Tet* KO compared to WT ExEpiLCs (X axis) versus the difference in 5mC levels between *Tet* KO and WT ExEpiLCs (Y axis). The dots are colored based on their peak cluster affiliation. Density distributions are plotted for each axis. **E.** Boxplots of 5mC levels in WT (blue) and *Tet* KO (grey) ExEpiLCs, for the peaks within the 5 clusters. **F.** Representative UCSC genome browser screenshot of M-ATAC and WGBS at a putative enhancer region (ENCODE cCREs) associated with the late opening cluster (C2). White lollipops represent unmethylated cytosines and black lollipops represent methylated cytosines. Blue shading indicates the putative enhancer region defined by our analysis. Methylation percentages of the highlighted region are indicated. *p < 0.05; **p < 0.01; ***p < 0.001; ****p < 0.0001; two-sided Wilcoxon test with Bonferroni correction.

ExEpiLCs showed the biggest number of differences, with 3071 peaks being more accessible and 3124 peaks being less accessible in *Dnmt* KO compared to WT (Figure 2B). Regions associated with chromatin closing in ExEpiLCs (transient and late closing clusters; C3 and C5, respectively) exhibited greater accessibility in *Dnmt* KO than in WT (Figure 2A,B), suggesting the requirement of 5mC to properly decommission this subset of candidate enhancers. Conversely, peaks with decreased accessibility were mostly found in the late opening cluster (C2; Figure 2B and Table S1). Gene ontology analysis revealed many terms associated with these regions were linked to gastrulation (Table S2). The decreased accessibility of late opening regions might reflect the inability of *Dnmt* KO cells to exit pluripotency^43,47^. When considering all non-promoter peaks, 7248 were more accessible and 9748 were less accessible in *Dnmt* KO compared to WT ExEpiLCs (Figure S2C), thus illustrating the extensive misregulation of the accessibility landscape in the absence of DNA methylation.

When taking 5mC ExEpiLC data into account, peaks with increased accessibility in *Dnmt* KO cells had higher endogenous 5mC than peaks with no differential accessibility, indicating the importance of 5mC for closing in WT conditions. Similarly, peaks with decreased accessibility in *Dnmt* KO had lower 5mC levels (Figure 2C and Table S3), suggesting a minor role of 5mC in regulating this class of regions. These trends were not observed at earlier time points, where 5mC levels are either low or in flux (Figure S2D). Together, these data suggest that the closing of chromatin during mESC-to-EpiLC differentiation at dynamic enhancer regions requires DNA methylation.

Surprisingly, only a minor fraction of statistically-significant differentially accessible regions was identified between *Tet* KO and WT ExEpiLCs (Figure S2E). Nevertheless, we observed a trend between a region’s accessibility and its cluster of origin. Opening peaks (C1 and C2) were overall less accessible in *Tet* KO, whereas closing peaks (C3, C4, and C5) showed increased accessibility compared to WT (Figure 2D). This lack of statistically-significant regions was observed despite higher levels of 5mC in *Tet* KO ExEpiLCs compared to WT, both at individual clusters (Figure 2D,E) and genome-wide (Figure S2F, S2G). Opening peaks had the starkest differences in 5mC, as illustrated by examples (Figure 2F, Figure S2H). These results suggest that while 5mC accumulation may impact a subset of enhancers, active demethylation by TET enzymes has an overall subtle effect on chromatin opening. Most regions ultimately opened despite a higher level of 5mC, thus uncoupling chromatin accessibility dynamics from 5mC removal during this stage of differentiation.

### M-CUT&Tag shows distinct epigenetic signatures of enhancer classes during pluripotency transitions

To characterize the epigenetic landscape of enhancers during the transition from naive to primed pluripotency, we employed M-CUT&Tag^28^ (Figure 3A) to simultaneously profile the enhancer-associated histone marks H3K4me1 or H3K27ac with DNA methylation across mESCs, EpiLCs, and ExEpiLCs. Results from WT, *Dnmt* KO, and *Tet* KO cells showed that opening enhancer regions gained both H3K4me1 and H3K27ac during differentiation, while closing regions lost these marks, consistent with trends observed in M-ATAC accessibility data (Figure 3B, Figure S3A-C, and Figure S4A-C).

**Figure 3.**
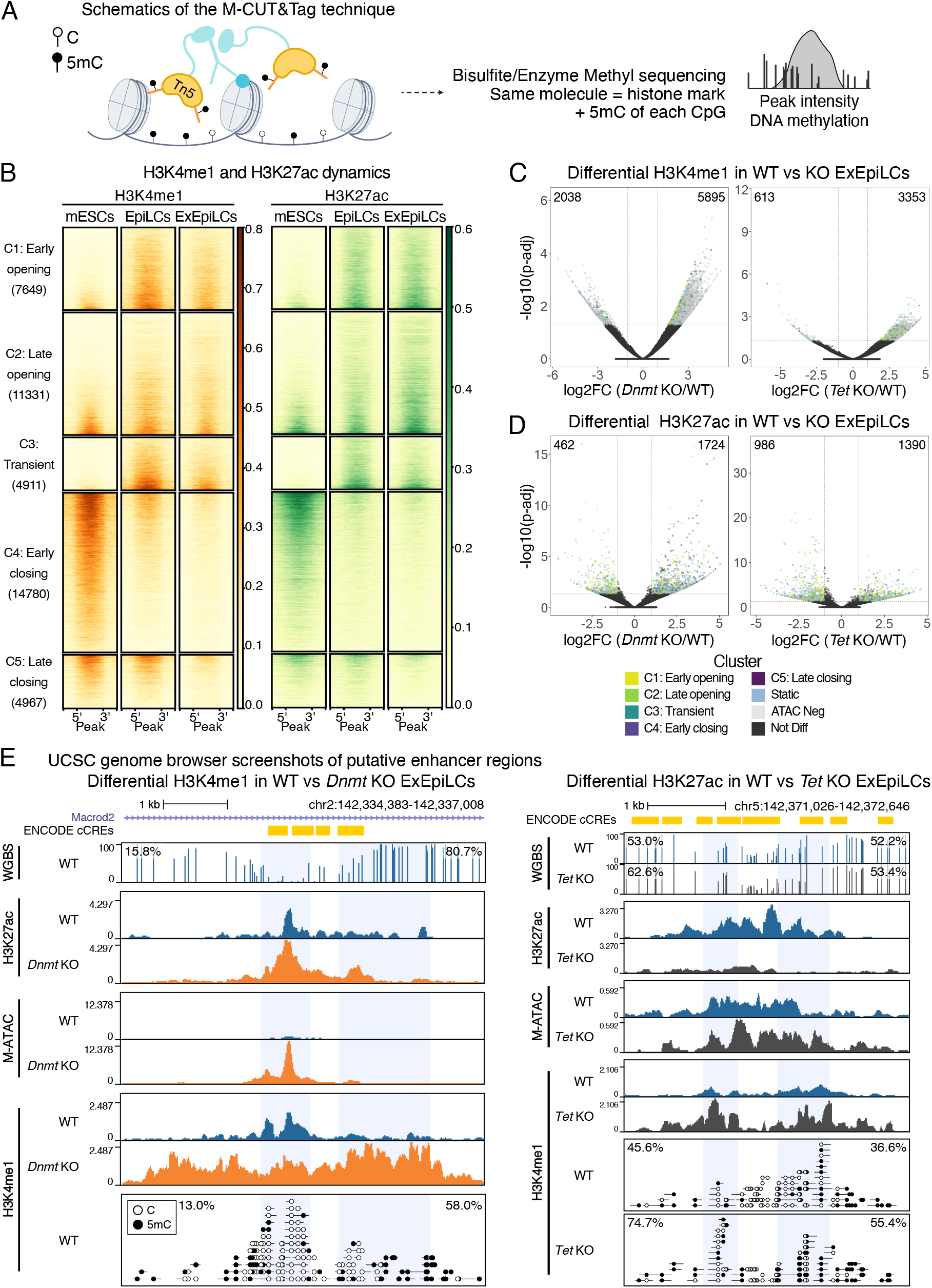
H3K4me1 and H3K27ac M-CUT&Tag reveal dynamic enhancer signatures. **A.** Schematic overview of the M- CUT&Tag technique. **B.** Heatmaps showing the H3K4me1 (left) and H3K27ac (right) M-CUT&Tag signal within the 5 clusters defined in Figure 1C. **C,D.** Volcano plots showing differential enrichment of H3K4me1 in WT vs *Dnmt* KO (left) or *Tet* KO (right) ExEpiLCs **(C)** and of H3K27ac in WT vs *Dnmt* KO (left) or *Tet* KO (right) ExEpiLCs **(D)** (log2-fold change > 1 or < -1; adjusted p-value < 0.05). M-CUT&Tag peaks are colored based on their overlap with ATAC-accessible regions (dynamic or static clusters) or lack thereof (ATAC Neg). **E.** Representative UCSC genome browser screenshots at putative enhancer regions (ENCODE cCREs, yellow boxes) with changing H3K4me1 in *Dnmt* KO ExEpiLCs and changing H3K27ac in *Tet* KO ExEpiLCs, compared to WT ExEpiLCs (left and right panels, respectively). White lollipops represent unmethylated cytosines and black lollipops represent methylated cytosines. Blue shading indicates the putative enhancer region defined by our analysis. Methylation percentages of the highlighted regions are indicated.

Differential enrichment analyses revealed heterogeneous changes in H3K4me1 signal for both KO cell lines, with gain and loss of histone mark levels across dynamic clusters, static peaks, and ATAC-negative regions (Figure 3C, Figure S3D, and Table S4). There was a relative gain in H3K4me1-enriched regions in *Dnmt* KO ExEpiLCs compared to WT (Figure 3C, left), concomitant with broadened H3K4me1 peaks (Figure 3E, Figure S3E,F) and in line with the perturbed accessibility landscape (Figure S2C). In *Tet* KO ExEpiLCs, the impact on H3K4me1 levels was less severe than in *Dnmt* KO (Figure 3C, right), with no differences to WT H3K4me1 levels in mESCs and EpiLCs (Figure S3D). These findings are consistent with the greater effect observed on the accessibility landscape in *Dnmt* KO relative to *Tet* KO during EpiLC differentiation, suggesting that *de novo* DNA methylation is required for establishing the enhancer landscape, while active demethylation has a more nuanced effect.

For the active mark H3K27ac, *Dnmt* KO and *Tet* KO displayed increased signal compared to WT mESCs across all ATAC clusters, with a prominent gain in early closing regions (C4) (Figure S4D and Table S4), suggesting this subset of enhancers might be particularly sensitive to changes in 5mC in the naive state. Substantial misregulation of H3K27ac was observed in *Tet* KO for all cell types (Figure 3D,E, Figure S4D, and Table S4). However, there was no clear bias towards decreased H3K27ac, which might have been expected in a DNA hypermethylated context. This apparent incongruence between extensive H3K27ac perturbations with only moderate H3K4me1 and accessibility changes in *Tet* KO will be discussed in a later section.

### H3K27ac M-CUT&Tag unveils relationship between enhancer activity and 5mC

We next classified putative enhancers based on the presence or absence of chromatin accessibility (M-ATAC), H3K4me1, and H3K27ac: active enhancers harbor the three features, primed enhancers are positive for H3K4me1 and chromatin accessibility, but negative for H3K27ac, and inaccessible enhancers are positive only for H3K4me1 (Figure 4A, Figure S5A). It should be noted that while H3K27ac is strongly associated with enhancer activation, its presence alone does not confirm activity, and further functional validation may be required.

**Figure 4.**
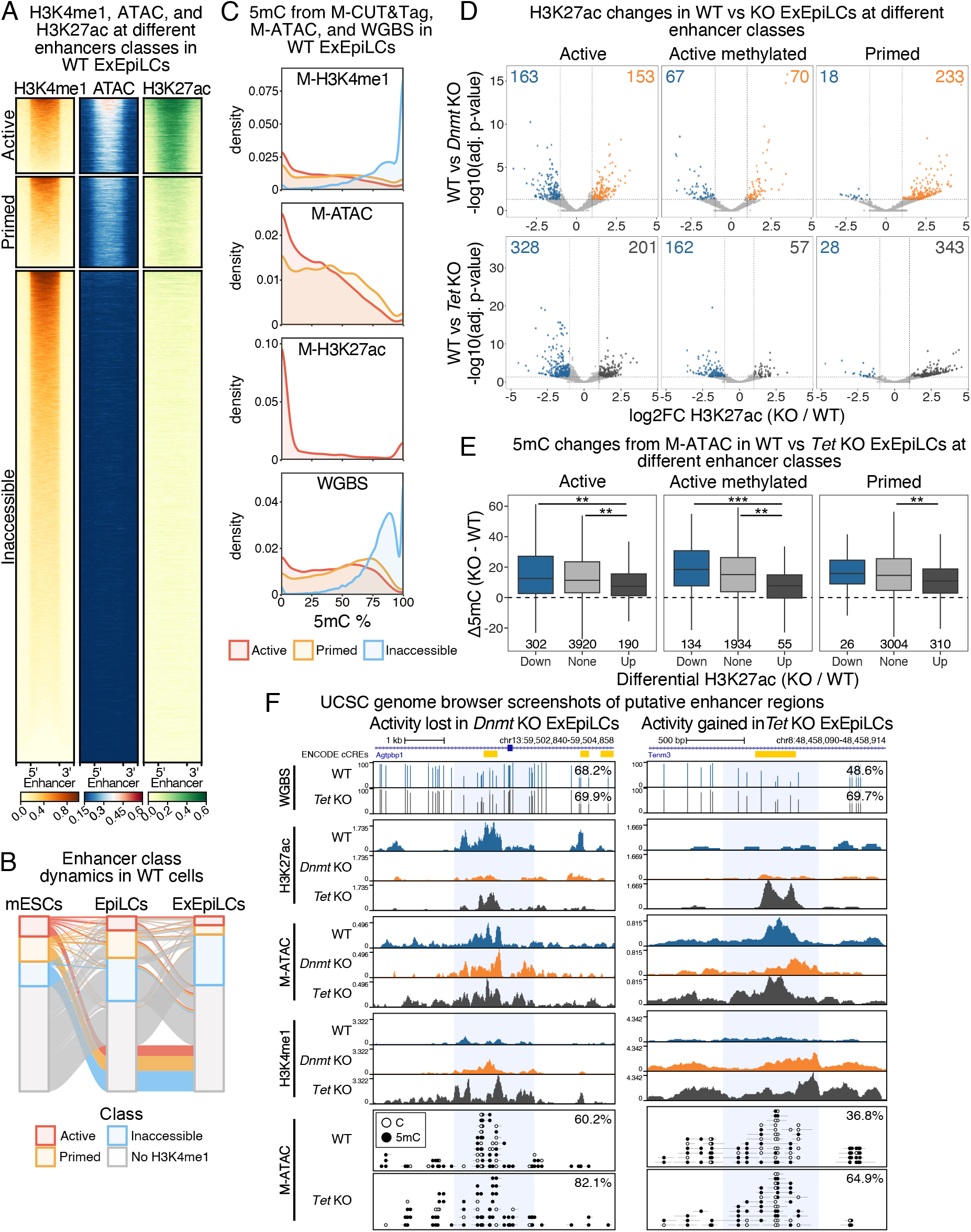
H3K27ac M-CUT&Tag reveals a complex interplay between enhancer activity and 5mC levels. **A.** Heatmaps showing the H3K4me1, M-ATAC, and H3K27ac signal in WT ExEpiLCs for enhancers classified as: active (positive for all three features), primed (positive for H3K4me1 and ATAC and negative for H3K27ac) and inaccessible enhancers (positive for H3K4me1 and negative for ATAC and H3K27ac). **B.** Alluvial plot representing the dynamics of the enhancer classes during mESCs to EpiLCs to ExEpiLCs differentiation in WT cells. Regions that lost the H3K4me1 signal in the corresponding day are depicted in grey. **C.** Density plots of 5mC levels at different classes of enhancers, quantified from M-ATAC, H3K4me1 and H3K27ac M-CUT&Tag, and WGBS datasets in WT ExEpiLCs. **D.** Volcano plots showing differential enrichment of H3K27ac in WT versus *Dnmt* KO (top) or *Tet* KO (bottom) ExEpiLCs at the different classes of enhancers (active, active methylated, and primed) (log2-fold change > 1 or < -1; adjusted p-value < 0.05). **E.** 5mC changes quantified from M-ATAC datasets at enhancers associated with gain or loss of H3K27ac in WT versus *Tet* KO ExEpiLCs, separated by enhancer class. The number of enhancers per box is indicated below. **F.** Representative UCSC genome browser screenshots of putative active methylated (left) and primed (right) enhancer regions (ENCODE cCREs). White lollipops represent unmethylated cytosines and black lollipops represent methylated cytosines. Blue shading indicates the putative enhancer region defined by our analysis. Methylation percentages of the highlighted regions are indicated. *p < 0.05; **p < 0.01; ***p < 0.001; ****p < 0.0001; two-sided Wilcoxon test with Bonferroni correction.

M-CUT&Tag and M-ATAC allowed us to highlight enhancer dynamics during differentiation from mESCs to EpiLCs to ExEpiLCs in WT cells (Figure 4B). Enhancers displayed considerable fluctuations in their activity state across time points, with (Ex)EpiLCs containing less active and more inaccessible enhancers than mESCs (Figure 4A,B and Figure S5A). An additional subset of regions lacked H3K4me1 signal in at least one of the three cell types, therefore losing the enhancer identity at a specific point of the mESC-to-EpiLC differentiation (Figure 4B and Figure S5B). Next, we quantified enhancer 5mC levels from M-ATAC, M-CUT&Tag (H3K4me1, H3K27ac), and WGBS datasets (Figure 4C, Figure S5C). As expected, 5mC levels were minimal in mESCs (Figure S5C). After differentiation into (Ex)EpiLCs the majority of enhancers gained 5mC, with the biggest effect observed at inaccessible regions (Figure 4C and Figure S5C). Contrary to the expected anticorrelation between H3K4me1 and 5mC^48^, we observed co-occurrence of both marks on the same molecule (Figure 3E, Figure 4C, Figure S3F, Figure S4E and Figure S5C). This observation held true for a subset of active enhancers (termed “active methylated”), which displayed 5mC levels of 50% or higher in M-ATAC and/or M-H3K4me1 datasets, suggesting that 5mC can coincide with an active enhancer state.

Both global loss and gain of 5mC (*Dnmt* KO and *Tet* KO, respectively) were associated with dysregulation of H3K27ac at all enhancer types (Figure 4D, Figure S6A). This included several primed enhancers that gained H3K27ac in mutant conditions. Changes in H3K27ac were accompanied by an increase of 5mC in *Tet* KO ExEpiLCs. While 5mC levels are overall higher in *Tet* KO than in WT cells (Figure S2G), enhancers with endogenous or decreased H3K27ac signal acquired more 5mC than regions with increased H3K27ac (Figure 4E, Table S5). This effect was limited to active enhancers in mESCs and EpiLCs (Figure S6B).

Our results suggest a complex interplay between H3K27ac and 5mC that goes beyond strict anticorrelation. Our data revealed 67 methylated enhancers that lost activity upon 5mC depletion (*Dnmt* KO), and 343 primed enhancers that became active in the absence of DNA demethylation (*Tet* KO), as exemplified in Figure 4F and Figure S6C. These challenges to the generally accepted 5mC-enhancer relationship underscore the nuanced, context-specific regulation occurring in the genome.

### hM-ATAC allows assessment of 5hmC dynamics at enhancers

Taken together, our results suggested that a subset of potentially active enhancers harbored substantial DNA methylation levels (≥ 50% 5mC) from M-ATAC and/or M-CUT&Tag. However, these techniques rely on the use of bisulfite conversion, which cannot discriminate between methylation and hydroxymethylation of cytosines (5hmC). Determining 5hmC levels is worthwhile *per se*, as 5hmC serves as an indicator of sites with active 5mC turnover, potentially leading to the recruitment of specific epigenetic readers^49^. To test 5hmC enrichment at enhancers, we developed hM-ATAC, which consists of the simultaneous detection of chromatin accessibility and 5hmC levels (Figure S7A), following the same principles as per M-ATAC with some notable differences (see Methods for details). Briefly, the cytosines on the modified oligonucleotides used for Tn5-associated adapters assembly and for oligonucleotide replacement harbored 5hmC instead of 5mC. After oligonucleotide replacement and gap repair, tagmented purified DNA was converted using the NEB 5hmC conversion module. This way, hydroxymethylated cytosines remained cytosines, while unmodified and methylated cytosines were converted to uracil and thymine, respectively. Following PCR and sequencing, 5hmC modifications could be detected similarly to 5mC in M-ATAC experiments.

We performed hM-ATAC in WT mESCs, EpiLCs, and ExEpiLCs. *Dnmt* KO and *Tet* KO ExEpiLCs—which should not harbor 5hmC—were used as controls. hM-ATAC displayed a strong Pearson correlation of ATAC signal with M-ATAC samples across cell types (mESCs: r = 0.77, EpiLCs: r = 0.81, and ExEpiLCs: r = 0.88, Figure S7B). 5hmC levels in hM-ATAC samples were substantially lower than 5mC quantified from M-ATAC (Figure 5A), in line with 5hmC corresponding to a fraction of the total DNA methylation estimated by bisulfite sequencing. As expected, *Dnmt* KO and *Tet* KO ExEpiLCs exhibited only background levels of 5hmC signal. By comparing our results with whole-genome 5hmC sequencing (5hmC-seq), hM-ATAC proved to be a more robust and cost-effective way to quantify 5hmC levels. We observed high agreement between both techniques for CpGs with at least ∼15x coverage in 5hmC-seq, whereas lower coverage resulted in inconsistent estimations (Figure S7C), emphasizing the strength of our targeted approach.

**Figure 5.**
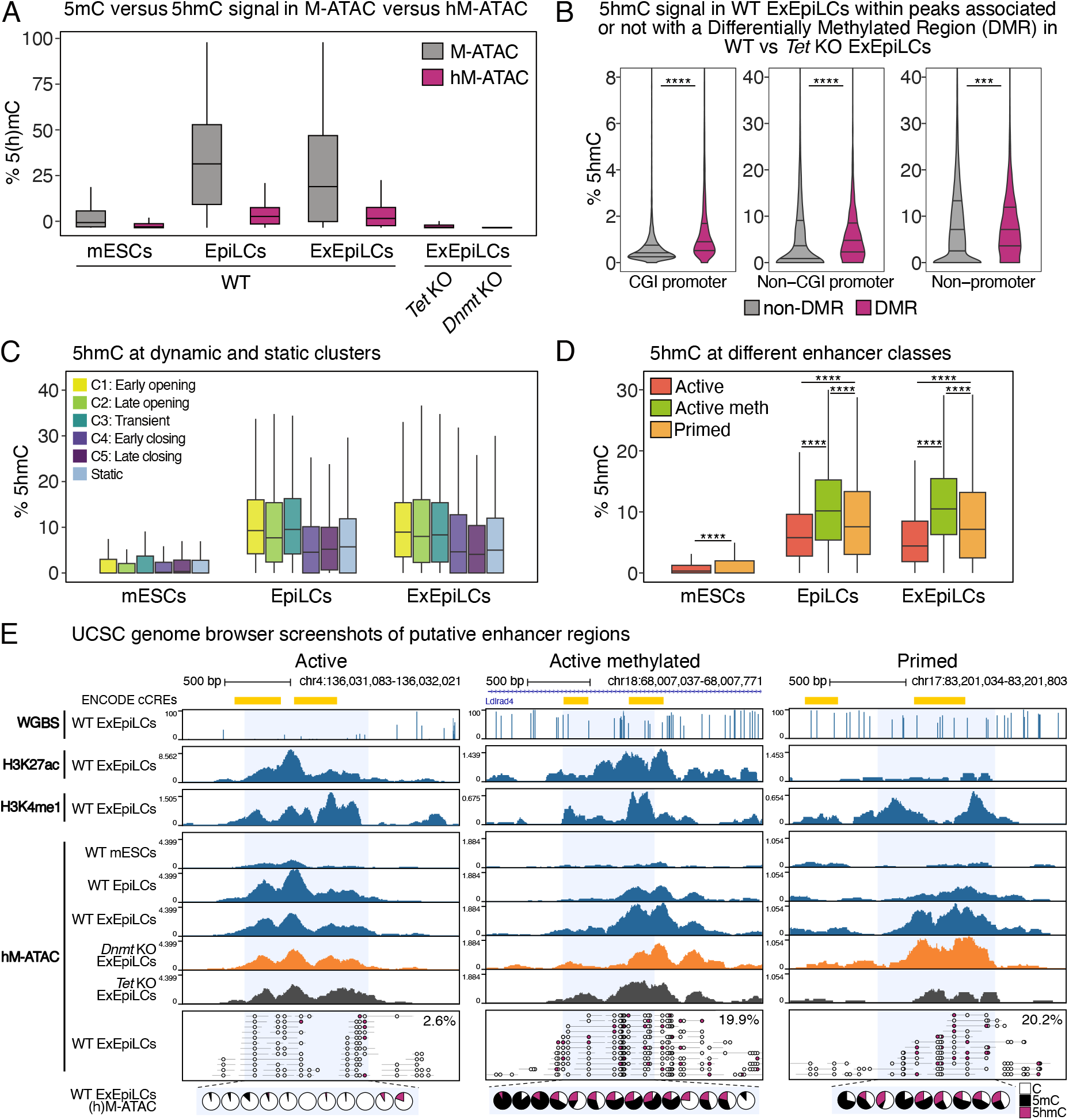
hM-ATAC assesses 5hmC dynamics at enhancers. **A.** Boxplots representing the levels of 5mC or 5hmC from M-ATAC (grey) versus hM-ATAC (magenta) samples. **B.** Violin plots representing the levels of 5hmC at hM-ATAC peaks, quantified in WT ExEpiLCs. Peaks were classified based on their genomic annotation (CGI promoter, non-CGI promoter, non-promoter) and further divided by their localization outside (grey) or inside (magenta) Differentially Methylated Regions (DMRs) in WT versus *Tet* KO ExEpiLCs (related to Figure S2F). **C.** Boxplots representing the mean levels of 5hmC from hM-ATAC in WT mESCs, EpiLCs, and ExEpiLCs for the five clusters of dynamic enhancers (related to Figure 1C) and static enhancers (related to Figure S1B). **D.** Boxplots representing the levels of 5hmC from hM-ATAC in WT mESCs, EpiLCs, and ExEpiLCs, comparing active enhancers (red), active methylated enhancers (green), and primed enhancers (orange). **E.** UCSC genome browser screenshots of putative enhancer regions (ENCODE cCREs). White and magenta lollipops represent unmethylated and hydroxymethylated cytosines, respectively. Blue shading indicates the putative enhancer region defined by our analysis. Hydroxymethylation percentages of the highlighted regions are indicated. The pie charts represent the proportion of C, 5mC, and 5hmC at each CpG in the shaded window, inferred from M- and hM-ATAC. *p < 0.05; **p < 0.01; ***p < 0.001; ****p < 0.0001; one-sided **(B)** or two-sided **(D)** Wilcoxon test.

Having successfully implemented the hM-ATAC method, we further assessed the distribution of 5hmC at different genomic locations. WGBS analysis identified 8858 differentially methylated regions (DMRs) with higher 5mC in *Tet* KO in comparison to WT ExEpiLCs (Figure S2F). As these regions are likely sites of TET activity, we compared 5hmC levels of hM-ATAC peaks within or outside DMRs. 5hmC was systematically higher at DMRs compared to matched regions, with non-promoter loci exhibiting the highest levels of 5hmC, followed by non-CGI and CGI promoters (Figure 5B, Table S6).

5hmC proved to be dynamic over time as well, as observed by the limited correlation of 5hmC levels between EpiLCs and ExEpiLCs (Figure S7D). This is likely a reflection of the dynamic nature of DNA methylation turnover during differentiation. Further evidence of fluctuating 5hmC patterns came from assessing 5hmC signatures at distinct enhancer types. Enhancers opening during the exit from naive pluripotency (clusters 1-3; see Figure 1C) showed increased 5hmC levels in (Ex)EpiLCs, whereas regions with stable or decreased accessibility (static peaks and clusters 4-5, respectively) showed only moderate 5hmC gains (Figure 5C). These patterns follow the opposite trend of 5mC levels at the same regions (Figure 1E), suggesting that while TET activity is not required for chromatin opening (Figure 2D and Figure S2E), active DNA demethylation co-occurs with increased chromatin accessibility. Overall, this indicates a functional role of TET enzymes to counter the wave of 5mC establishment genome wide, and contribute to the epigenetic landscape of enhancers that are being licensed.

Next, we analyzed the relationship between enhancer class and 5hmC. Both active and primed enhancers displayed higher 5hmC signals in differentiated cells compared to mESCs, with only minimal differences in 5hmC levels between EpiLCs and ExEpiLCs (active methylated enhancers are not found in mESCs). Notably, active methylated enhancers harbored higher 5hmC levels than the remainder active subset in (Ex)EpiLCs, while primed enhancers exhibited intermediate levels of 5hmC (Figure 5D, Table S6). This may suggest that a large proportion of the active methylated class are on a trajectory towards lower methylation levels, which may appear at further differentiation stages. Examples are shown of active, active methylated, and primed enhancers (Figure 5E and Figure S7E).

In summary, hM-ATAC is not only an affordable and easy-to-implement technique to assess 5hmC dynamics at regulatory elements, but allows for novel insights into TET-mediated enhancer regulation.

### Enhancers altered in DNA methylation mutants impact gene expression

In order to investigate the relationship between enhancer class and target gene expression, we performed RNA-seq in *Tet* KO mESCs, EpiLCs, and ExEpiLCs and analyzed them alongside already published WT and *Dnmt* KO datasets in the corresponding cell types^43^. Principal Component Analysis (PCA) revealed a globally coherent trajectory from mESCs to EpiLCs to ExEpiLCs for the three genotypes (Figure 6A). Our RNA-seq results may also help explain the unexpected delay in 5mC accumulation observed in *Tet* KO EpiLCs compared to WT (Figure S2G), as *Dnmt3a* and *Dnmt3l* expression peaks at EpiLCs were substantially lower in *Tet* KO cells (Figure S8A).

**Figure 6.**
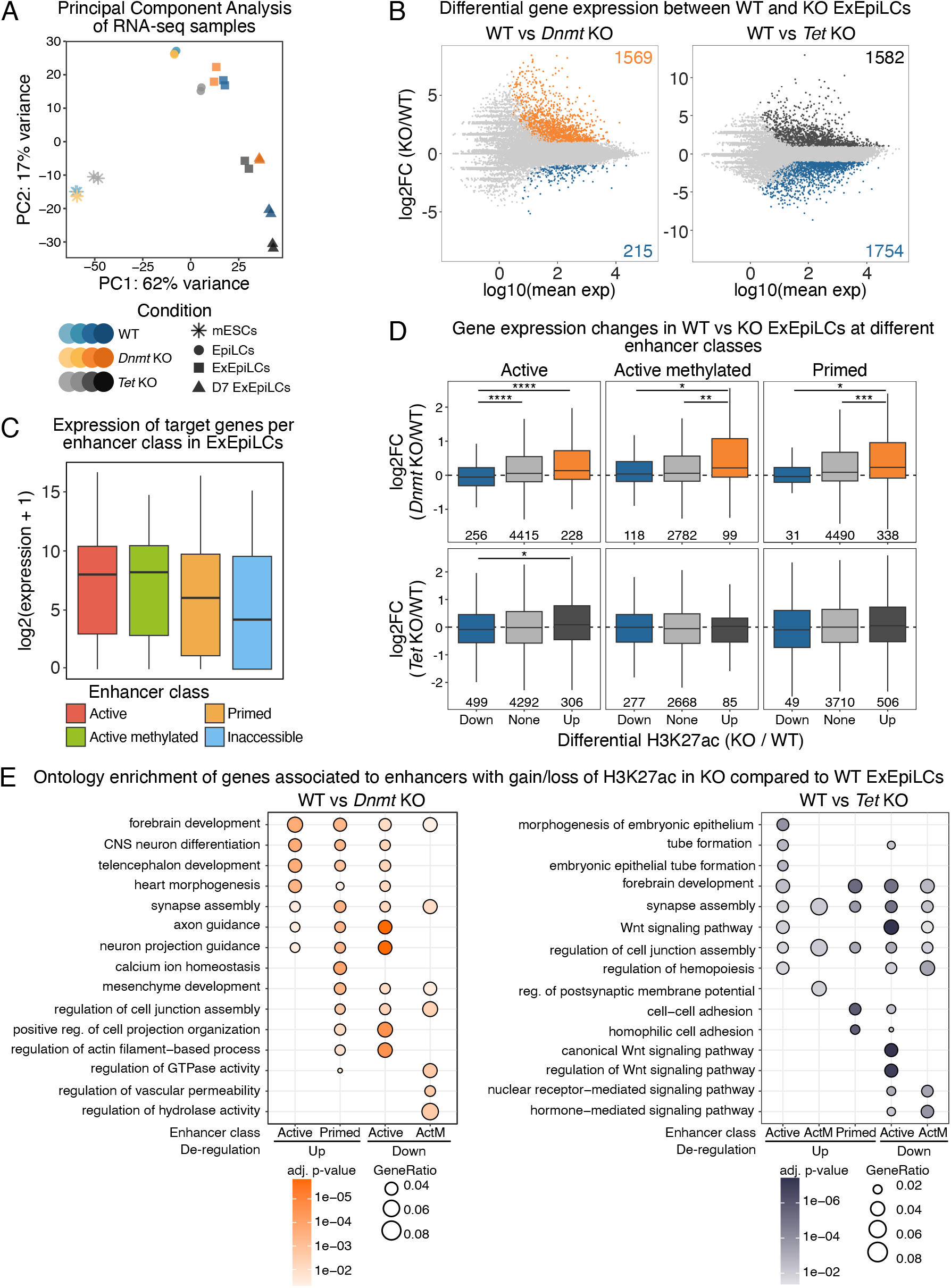
Enhancer state and DNA methylation influence gene expression in a context-dependent manner. **A.** Principal Component Analysis (PCA) of RNA-seq samples for WT and KO across differentiation. **B.** MA plots of differential gene expression in ExEpiLCs of WT vs *Dnmt* KO (left), and WT vs *Tet* KO (right). Number of up-regulated and down-regulated genes is indicated. **C.** Normalized read counts of genes associated with each enhancer class in WT ExEpiLCs. Enhancer-gene association was performed with GREAT (see Methods). **D.** RNA expression differences in genes associated with enhancers displaying gain/loss of H3K27ac in WT vs KO ExEpiLCs, separated by enhancer class. The number of genes per box is indicated below. **E.** Ontology enrichment of genes associated with enhancers with gain/loss of H3K27ac in *Dnmt* KO (left) or *Tet* KO (right) compared to WT ExEpiLCs, separated by enhancer class. ActM = active methylated. *p < 0.05; **p < 0.01; ***p < 0.001; ****p < 0.0001; two-sided Wilcoxon test with Bonferroni correction.

*Tet* KO displayed an apparent accelerated differentiation phenotype compared to WT and *Dnmt* KO cells. This observation is consistent with greater gene expression changes—both in terms of up-regulation and down-regulation—in *Tet* KO than in *Dnmt* KO when compared to WT mESCs and EpiLCs (Figure S8B). It might also clarify the substantial number of differential H3K27ac peaks between *Tet* KO and WT (Ex)EpiLCs (Figure 3D and Figure S4D). In ExEpiLCs after 4 or 7 days of differentiation, we observed increased gene up-regulation in *Dnmt* KO cells compared to WT (Figure 6B and Figure S8B). Expression of puripotency and differentiation markers is comparable between the three genotypes with some exceptions, such as elevated levels of the EpiLC marker *Oct6* in *Tet* KO EpiLCs (Figure S8C), in agreement with the accelerated phenotype observed in the PCA (Figure 6A).

Enhancers from previously identified classes were linked to potential target genes using Genomic Regions Enrichment of Annotations Tool (GREAT^50^). As expected, genes associated with active enhancers (including active methylated) were more highly expressed than genes associated with primed or inactive enhancers (Figure 6C and Figure S8D). Gain of H3K27ac in *Dnmt* KO ExEpiLCs (and, to a lesser extent, *Tet* KO) was concomitant with increased gene expression of target genes (Figure 6D, Table S7). This effect was less prominent in mESCs and EpiLCs (Figure S9A,B), indicating a more minor role for DNA (de)methylation machinery on gene regulation early in differentiation (Figure 4E, Figure S6B). Levels of 5mC/5hmC at primed enhancers (no H3K27ac) showed no direct impact on gene expression, whereas a mild correlation could be observed for active enhancers (more 5mC/5hmC, less expression; Figure S9C). Our results suggest a complex interplay between 5mC/5hmC and enhancer activity. Enhancers with altered H3K27ac levels in *Dnmt* KO and *Tet* KO were associated with genes linked to differentiation terms that occur after gastrulation (Figure 6E), exemplifying that proper 5mC deposition and removal at enhancers is likely important for priming pluripotent cells for germ layer specification.

### (h)M-ATAC reveals 5mC-responsive transcription factor regulation

As mentioned above, gene expression regulation by enhancers is driven in part by the ability of TFs to bind to their motifs. Therefore, we investigated whether specific TFs, or TF classes, were linked to chromatin accessibility and enhancer state, as well as their association with 5mC.

A total of 805 TF motifs were significantly enriched within the 5 clusters that represent chromatin accessibility dynamics in WT cells, agnostic to the presence of a CpG site in the motif (Figure S10A, Table S8). Clusters C4 and C5 (early and late closing, respectively) had comparable motif enrichment profiles, indicating a shared subset of TFs involved in chromatin closing. Conversely, clusters C1 and C3 (early opening and transient) had profiles more similar to each other than to C2 (late opening), which displayed an overall depletion of transcription factor binding sites. Our results suggest that during the mESC-to-EpiLC differentiation, the timing of chromatin opening (rather than chromatin opening, *per se*) has a stronger link to motif enrichment, despite transient regions closing at further developmental stages.

Next, we looked for motifs driving differential accessibility in WT versus *Dnmt* KO cells. We also incorporated information for TF 5mC-specificity or -sensitivity based on published MethylSelex data^5^ when available. Both CpG- and non-CpG-containing motifs were found enriched at differentially accessible regions for every cell type (Figure S10B), with a large proportion of motifs discovered within peaks exhibiting increased chromatin accessibility in *Dnmt* KO (Ex)EpiLCs (Figure S10B; Figure 7A, orange dots). Notably, several of these motifs have been linked with TFs that are responsive to 5mC presence (Figure 7A, bottom panel, red, blue, and green dots). As a general trend, we observed that within peaks that are more accessible in *Dnmt* KO ExEpiLCs, there is an enrichment of motifs in which TFs exhibit preferential binding to 5mC (MethylPlus) (Figure 7B, peaks up; Table S9).

**Figure 7.**
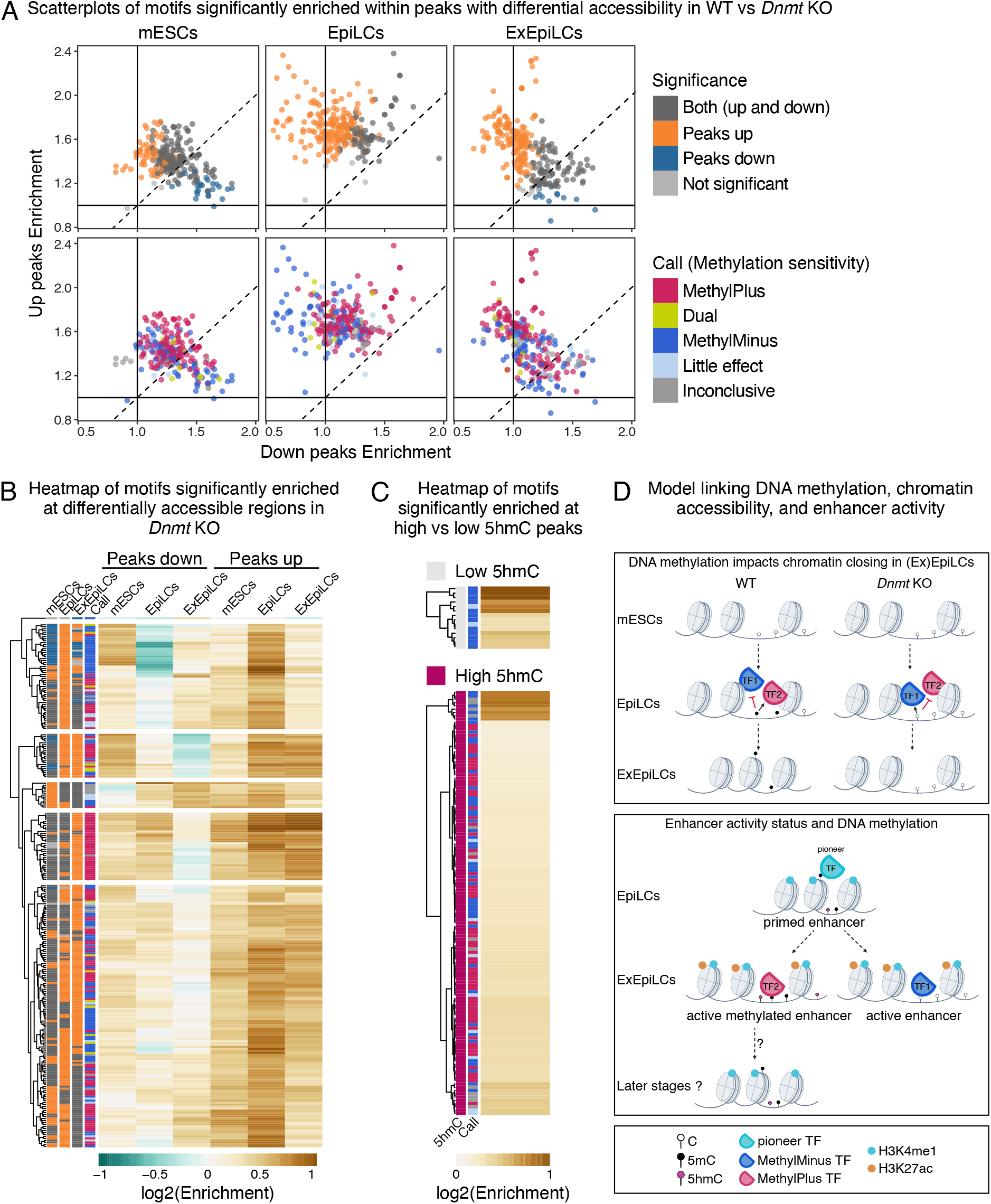
Data reveals 5mC-responsive transcription factor regulation. **A.** Scatterplots of motifs significantly enriched at peaks with differential accessibility in WT vs *Dnmt* KO. The motif enrichment score (see Methods) in down peaks (accessibility lost in *Dnmt* KO; X axis) and up peaks (accessibility gained in *Dnmt* KO; Y axis) is compared for each cell type. Motifs are colored based on their enrichment in up or down peaks (top), or their methylation sensitivity obtained from MethylSelex (Yin et al., 2017) (bottom). The dotted line represents x = y. **B.** Enrichment score heatmap of motifs represented in **A**. Peak group enrichment and methylation sensitivity are depicted on the left. Only motifs with characterized methylation sensitivity are shown, see Figure S10B for a heatmap including all motifs. **C.** Heatmap of motifs significantly enriched at high versus low 5hmC hM-ATAC peaks in ExEpiLCs. Color codes are shared through panels **A-C**. **D.** Model linking DNA methylation, chromatin accessibility, and enhancer activity. Top: DNA methylation impacts chromatin closing in (Ex)EpiLCs in two ways. Some TFs will bind preferentially to unmethylated CpGs (MethylMinus). In EpiLCs, these TFs would bind *Dnmt* KO-specific accessible regions at a higher rate. Alternatively, other TFs will bind preferentially to methylated CpGs (MethylPlus) and will be required for chromatin closing in EpiLCs. In *Dnmt* KO cells, such TFs would bind less efficiently and the chromatin would stay accessible. Bottom: Proposed dynamics of methylated enhancers. For enhancers primed in EpiLCs and active in ExEpiLCs, half of them will harbor high methylation levels (≥ 50% 5mC) and also elevated 5hmC levels. These enhancers will be bound by a different set of TFs than the active lowly methylated enhancers (MethylPlus versus MethylMinus TFs, respectively). MethylPlus TFs may either contribute to the active state or facilitate enhancer decommissioning at later developmental stages.

Motif enrichment also distinguished regions of high (> 17% 5hmC, top decile) versus low (< 0.2% 5hmC, bottom decile) 5hmC levels. Motifs enriched at high 5hmC regions included similar proportions of CpG- and non-CpG binders, whereas regions with low 5hmC were enriched almost exclusively in CG-containing motifs (Figure S10C and Table S10). These motifs corresponded to TFs that prefer unmethylated cytosines (MethylMinus). At regions with a high 5hmC signal, motifs trended towards MethylPlus TFs (Figure 7C and Figure S10C), bearing in mind high 5mC and 5hmC levels are correlated at primed and active methylated enhancers (Figure 5D). Taken together, we can refine a model of the intricate interplay between TFs and 5mC dynamics, which we elaborate on below (Figure 7D, see Discussion).

## Discussion

The integration of M-ATAC, H3K27ac/H3K4me1 M-CUT&Tag, and hM-ATAC has provided a high-resolution, multi-layered view of the epigenetic reprogramming of enhancers during the transition from naive to primed pluripotency. These findings not only deepen our understanding of enhancer dynamics but also challenge traditional views of DNA methylation as a purely repressive mark.

The observation that chromatin accessibility and 5mC can coexist at enhancers during the transition from mESCs to EpiLCs and ExEpiLCs suggests that DNA methylation does not universally correlate with chromatin closure. Instead, 5mC may play a nuanced role in fine-tuning enhancer activity, potentially by modulating transcription factor binding or stabilizing chromatin states. This highlights the need for further investigation into the context-dependent roles of 5mC at enhancers. The requirement of DNA methylation for chromatin closing during differentiation, as demonstrated in *Dnmt* KO cells, underscores the importance of 5mC in establishing a repressive chromatin environment. However, the fact that TET-mediated demethylation is not required for chromatin opening suggests that other mechanisms, such as pioneer transcription and/or chromatin remodeling factors, may drive enhancer activation independently of 5mC removal, or that TET may have more important functions in future developmental stages.

H3K4me1 is widely regarded as a general enhancer mark, present at both active and primed enhancers regardless of their transcriptional state^1,2^. Unlike H3K27ac, which is dynamically regulated during differentiation, H3K4me1 is considered as reflecting the potential rather than the activity of enhancers^51^. In *Tet* KO cells, H3K4me1 levels remained largely unchanged compared to WT. This stability aligns with the conventional view of H3K4me1 as a constitutive enhancer mark. In *Dnmt* KO cells, however, we observed broadened H3K4me1 peaks and mixed increases and decreases in H3K4me1 signal across dynamic clusters, static regions, and even ATAC-negative regions. The broadening of H3K4me1 peaks and the global misregulation in H3K4me1 signal in *Dnmt* KO cells suggest that DNA methylation could play a role in defining the boundaries and stability of enhancer regions. The coexistence of 5mC and H3K4me1 may reflect a primed enhancer state, where enhancers are marked for potential activation until specific developmental cues are received. Alternatively, 5mC at primed enhancers could represent a transient state that will erode when activated, which is consistent with elevated levels of 5hmC observed at primed enhancers compared to the active unmethylated class (Figure 5D).

The discovery that a subset of enhancers harbor active chromatin features (ATAC+, H3K4me1+, H3K27ac+) alongside elevated 5mC levels (≥ 50%) is particularly striking. This suggests that 5mC may not always act as a repressor but could instead positively contribute to the enhancer activity, aligning with previous evidence of "bivalent" enhancers^14^, where repressive and active marks coexist, allowing for rapid transitions between transcriptional states. Alternatively, we could imagine that 5mC might help maintain enhancer specificity by modulating the binding of specific readers that will further be required for closing the chromatin at later developmental stages. Our motif analysis supports the latter point, as we observed enrichment of MethylPlus TF motifs in *Dnmt* KO peaks in ExEpiLCs. We propose a model in which MethylPlus TFs are required for decommissioning enhancers in WT; in *Dnmt* KO, the lack of 5mC results in persistent ATAC positive regions (Figure 7D, top). This is not mutually exclusive from the more traditional models of MethylMinus TFs regulating lowly methylated enhancers^52,53^, but adds a layer of nuance for the mechanism of chromatin closing. Future work will be necessary to interrogate this model.

It is reasonable to ask if MethylPlus TFs contribute to the high proportion of active methylated enhancers that we described. In other words, is it possible that there is a large fraction of enhancers that require 5mC for their activity that have been overlooked? If this were the case, we would expect that many enhancers within the active methylated class would lose the H3K27ac mark in *Dnmt* KO. While we did find such examples (Figure 4F and Figure S6C), these were very small in number. Although globally our data does not suggest that 5mC, *per se*, plays a major role in activation, it will be worthwhile to formally test candidate active methylated enhancers for their 5mC requirements with tools such as epigenome editing. It is notable that active methylated enhancers also exhibit high levels of 5hmC (Figure 5D). This suggests that they are TET targets, and on a trajectory towards demethylation. We propose that the majority of active methylated enhancers represent a transient state, and are bound by TFs that either are MethylPlus, or are simply agnostic to 5mC levels (i.e., do not have a CpG in their binding motif) (Figure 7D, bottom). Continued differentiation to various lineages would determine whether the 5mC and/or enhancer activity state changes.

The mESC-to-EpiLC differentiation system represents a developmental window that ends at primed pluripotency, just prior to germ layer specification. Our findings suggest that 5mC could be involved in "priming" enhancers for future activation, particularly in developmental contexts where precise timing of gene expression is critical. Our hM-ATAC experiments have shown that 5hmC could also be a mark of such primed enhancers (Figure 5D), consistent with a recent study^54^. As technology evolves, future work will allow for analysis of 5mC and 5hmC on the same molecule^54^. Both DNMT and TET enzymes are required in somatic cell types, thus it is possible 5mC and 5hmC plays a more important role as enhancer elements in more differentiated states. The primed enhancers we uncovered in ExEpiLCs would presumably be activated in specific lineages, and potentially misregulated in 5mC mutants. With the emergence of sophisticated differentiation techniques^53^ and PROTAC-mediated acute degradation of essential proteins, hopefully these aspects will be assessed in the coming years.

### Limitations

While our study provides novel insights into the interplay between 5mC and enhancer activity, our analysis focused primarily on H3K27ac as a marker of enhancer activation^2^. However, other histone acetylation marks, such as H3K9ac or H3K16ac, may also contribute to enhancer activity and were not examined in this study^55–57^. Additionally, we observed correlations between 5mC, chromatin accessibility, and histone modifications, but causal relationships remain to be established. Functional perturbation experiments, such as CRISPR-mediated targeted manipulation of DNA methylation, would be necessary to confirm the regulatory roles of 5mC at enhancers.

## Acknowledgments

We thank Anaëlle Azogui, Sarah Battault, and Elisa Carretin for experimental assistance. We thank members of the Greenberg lab for useful discussions. All sequencing was performed by Novogene, Co., Ltd. This work was supported by the European Research Council (ERC-StG-2019 DyNAmecs), funds from the Agence National de Recherche (ANR, projects ANR-21-CE12-0015-03, REMEDY and ANR-25-CE12-6449, MethylEnh) and the Fondation CNRS Georges Brahms Prize awarded to P.L. J.R.A was supported by a Fondation pour la Recherche Médicale, Post doc France Fellowship (SPF202110014238); A.D is supported by a 4th year PhD fellowship from the Fondation ARC.

## Author contributions

P.L. and M.V.C.G. were responsible for planning the experimental design. P.L., B.D., P.A.R., H.B. and A.D. performed the experiments. M.M.-F., J.R.A., A.B.B. and A.F.B. were responsible for data analysis, data visualization, statistics, and computational approaches, with assistance from P.L., B.D., and P.A.R. M.V.C.G., P.L. and P.-A.D. acquired funding. M.M.-F., P.L. and M.V.C.G. wrote and assembled the manuscript.

## Data and code availability

Genomic data has been uploaded to the Gene Expression Omnibus (GEO) under accession numbers GSE336120, GSE336122, GSE336124, GSE336125, GSE336126, and GSE336127. Scripts for library processing can be found at https://github.com/julienrichardalbert/sra2bw. Scripts for data analysis and visualization can be found at https://github.com/marletmorales/MethylEnh.

## Methods

### Cell lines

E14Tg2a (E14) mouse ESCs was the parental line used for all experiments in this study. Both the *Dnmt* KO and *Tet* KO were generated in-house using CRISPR/Cas9, and described previously^58,59^.

### Cell culture

Cells were cultured as previously described^60,61^. Briefly, The ESCs were grown on 0.1% gelatin-coated flask in an incubator at 37°C and 5% CO2, in serum culture conditions using Glasgow medium (Gibco) supplemented with 15% Fetal bovine Serum (FBS), 0.1 mM MEM non-essential amino acid, 1 mM sodium pyruvate, 2 mM l-glutamine, penicillin, streptomycin, 0.1 mM β-mercaptoethanol and 1000 U/ml leukemia inhibitory factor (LIF), and passed with trypsin. Before EpiLC differentiation, cells were transitioned from serum to 2i + Vitamin C (vitC) cell culture conditions for at least 7 days. For the 2i + vitC culture conditions, N2B27 medium (50% neurobasal medium, 50% DMEM) supplemented with N2 (Gibco), B27 (Gibco), 2 mM l-glutamine, 0,1 mM β-mercaptoethanol, penicillin, streptomycin, LIF and 2i (3 μM Gsk3 inhibitor CT-99021, 1 μM MEK inhibitor PD0325901) and ascorbic acid (vitC, Sigma) at a final concentration of 100 μg/ml. Cells were passed using Accutase (Gibco) every 2 days. The transition from serum-grown high DNA methylation cells to 2i+VitC-grown low DNA methylation levels was assessed using LUminometric Methylation Assay (LUMA)^62^.

To induce EpiLC differentiation, cells were replated at a density of 2 × 10^5^ cells/cm2 on Fibronectin (10 μg/ml, Sigma))-coated plates in N2B27 medium supplemented with 12 ng/ml FGF2 (R&D) and 20 ng/ml Activin A (R&D) and passed with Accutase at day 3 of differentiation for cells harvested at day 4 or day 7.

### M-ATAC and M-CUT&Tag

#### Transposase assembly

The Tn5 transposome was assembled with methylated adaptors as per the T-WGBS protocol^63^. (Tn5mC-Apt1 and Tn5mC1.1-A1block; 100 μM each; Table S11) and annealed in a thermomixer with the following program: 95 °C for 3 min, 70 °C for 3 min, 45 cycles of 30 s with a ramp at − 1 °C per cycle to reach 26 °C. Eight microliters of annealed adapters were incubated with 50 μl of house-made pA-Tn5 transposase^60^ at room temperature for 60 min to assemble the transposome and stored at -20 °C for future use.

#### M-ATAC

M-ATAC was performed as previously reported^24,25^. Briefly, M-ATAC was performed with 50,000 cells as per the original ATAC-seq protocol^64^. Cells were washed in cold PBS and resuspended in 50 μl of cold lysis buffer (10 mM Tris-HCl, pH 7.4, 10 mM NaCl, 3 mM MgCl2, 0.1% IGEPAL CA-630). The tagmentation reaction was performed in 25 μl of freshly prepared tagmentation buffer (10 mM Tris-HCl pH 8.0, 5 mM MgCl2 and 10% v/v dimethylformamide), 2.5 μl transposase containing the methylated adaptors (see section “Transposase Assembly” for details), and 22.5 μl of nuclease-free H2O at 37 °C for 30 min. Purified DNA (on column with the Qiagen Mini Elute kit) was bisulfite converted (see section “Bisulfite conversion” for details) and PCR amplified (see “Amplification of M-ATAC-seq and M-C&T libraries” for details).

#### M-CUT&Tag

M-CUT&Tag was performed as previously reported^28^ with some modifications. Briefly, CUT&Tag of H3K4me1 and H3K27ac was performed using pA-Tn5 loaded with a methylated adapter, following the bench-top CUT&Tag protocol^29^. H3K4me1-CUT&Tag was conducted with 500,000 cells, while H3K27ac-CUT&Tag was conducted with 1 million cells. Harvested cells were counted with an automated cell counter.

The cell pellet was washed twice with 1 ml Wash Buffer (20 mM HEPES-KOH pH 7.5, 150 mM NaCl, 0.5 mM Spermidine, 1× Protease inhibitor cocktail), and then resuspended in 200 μL of Wash Buffer. For H3K27ac CUT&Tag, nuclei were extracted as previously^65^ by incubating cells for 10 minutes on ice in 200 µL/sample of cold Nuclei Extraction buffer (NE buffer: 20 mM HEPES-KOH pH 7.9, 10 mM KCl, 0.1% Triton X-100, 20% Glycerol, 0.5 mM Spermidine, 1x Protease Inhibitor cocktail).

Following incubation in the NE buffer, nuclei were centrifuged for 3 min at 600g at RT, then resuspended in 100 µL cold NE buffer. For the cells or nuclei to bind to Concanavalin A-coated magnetic beads, 10 μL activated beads were added into each sample and the cell-bead mixture was rotated at RT for 10 min. The tubes were then placed on a magnet stand to clear and the supernatant was removed. The bead-bound cells were resuspended in 100 μL of Antibody Buffer (20 mM HEPES pH 7.5, 150 mM NaCl, 0.5 mM Spermidine, 1× Protease inhibitor cocktail, 0.01% Digitonin, 2 mM EDTA, 0.1% BSA) with anti-H3K4me1 antibody (Cell Signaling Technology 5326), or anti-H3K27ac antibody (active motif 39133) at 1:100 dilution. Primary antibody incubation was performed at 4°C overnight on a rotator. The tubes were then placed on the magnet stand and the supernatant was discarded. Next, the bead-bound cells were resuspended in 100 μL of Antibody Buffer with Guinea Pig anti-Rabbit IgG antibody at 1:100 dilution and then incubated at RT for 30 min on a rotator. After secondary antibody binding, the cells on beads were washed three times with 1 mL Dig-Wash Buffer (20 mM HEPES pH 7.5, 150 mM NaCl, 0.5 mM Spermidine, 1× Protease inhibitor cocktail, 0.01% Digitonin) The bead-bound cells were then incubated with pA-Tn5 loaded with methylated adapter at 1:200 dilution in either 100 μL Dig-300 Buffer (20 mM HEPES pH 7.5, 300 mM NaCl, 0.5 mM Spermidine, 1× Protease inhibitor cocktail, 0.01% Digitonin) for H3K4me1 CUT&Tag, or Dig-Med buffer for H3K27ac CUT&Tag (20 mM HEPES pH 7.5, 300 mM NaCl, 0.5 mM Spermidine, 1× Protease inhibitor cocktail, 0.05% Digitonin) for 1 hour at RT on a rotator. After pA-Tn5 binding, the cells on beads were washed three times with 1 mL Dig-300 Buffer to remove unbound pA-Tn5. To activate pA-Tn5 for tagmentation, the bead-bound cells were resuspended in 100 μL Tagmentation Buffer (20 mM HEPES pH 7.5, 300 mM NaCl, 0.5 mM Spermidine, 1× Protease inhibitor cocktail, 0.01% Digitonin, 10 mM MgCl2) then incubated at 37°C for 1 hour in a thermomixer with shaking at 1000 rpm. For H3K4me1 CUT&Tag, the tagmentation reaction was purified using the MinElute kit (Qiagen) according to the manufacturer’s instructions. For H3K27ac, 2.25 µL of 0.5 M EDTA, 2.75 µL of 10% SDS and 0.5 µL of 20 mg/mL Proteinase K was added to 50 µL of sample To stop tagmentation, which was incubated at 55 °C for 30 min or overnight at 37 °C, and then at 70 °C for 20 min to inactivate Proteinase K. DNA was extracted using a Phenol:Chloroform:Isoamyl Alcohol solution.

Oligonucleotide replacement and gap repair was performed as previously reported^24,25^. Briefly, 11μL M-ATAC/M-CUT&Tag DNA eluate, 2μL 10μM Tn5mC-Repl01 oligo, 2μL 10x ampligase buffer, 2μL dNTPs 2.5mM each was assembled in a PCR tube and incubated as follows in a PCR thermocycler: 50°C for 1 minute, 45°C for 10 minutes, ramp down to 37°C at a rate of −0.1°C/second, hold at 37°C. Then 1μL T4 DNA polymerase and 2.5μL ampligase were added without removing the tube from the thermocycler and the reaction was incubated at 37°C for 30 minutes, then held at 4°C before either bisulfite conversion or EM-seq.

#### Bisulfite conversion of M-ATAC and H3K4me1 M-CUT&Tag DNA

For M-ATAC and H3K4me1 M-CUT&Tag, DNA material following oligonucleotide replacement and gap repair was bisulfite converted according to manufacturer instructions using the Zymo EZ DNA Methylation-Gold Kit (catalog number D5006).

#### Enzymatic Methyl-seq of H3K27ac M-CUT&Tag DNA

For H3K27ac M-CUT&Tag, DNA material following oligonucleotide replacement and gap repair steps (see previous sections) was converted using the NEB Enzymatic-Methyl-seq kit as per the manufacturer instructions.

#### WGBS

Genomic DNA was extracted using NucleoSpin tissue kit (Macherey-Nagel ref. 740952) as per the manufacturer instructions. DNA was treated with RNAse-A (1mg/ml final) at 37°C for 30 minutes and then Proteinase K (0.5mg/ml final) at 70°C for 15 minutes. DNA was re-purified using phenol:chloroform:isoamyl-alcohol and ethanol precipitated and was sent to novogene for library preparation using the Accel-NGS Methyl-Seq DNA library kit (Swift Biosciences) and sequencing using 2x150bp paired-end reads on the NovaSeq6000 instrument.

#### Detection of 5hmC

5hmC was detected either from M-ATAC material (hM-ATAC) or from genomic DNA for genome-wide control (5hmC-seq).

hMATAC was performed as per M-ATAC (see M-ATAC section above) with the following modifications: the transposase has been assembled with hydroxymethylated cytosines (IDT), and the tagmented purified DNA has been processed through the NEB Enzymatic 5hmC-seq Conversion Module kit as per the manufacturer instructions.

For 5hmC-seq, genomic DNA was extracted using NucleoSpin tissue kit (Macherey-Nagel ref. 740952) as per the manufacturer instructions and sonicated using covaris instrument. Sonicated DNA was processed through the NEB Enzymatic 5hmC-seq complete kit as per the manufacturer instructions.

#### PCR amplification

DNA was amplified and barcoded in 30μL PCR reactions (15μL 2x KAPA HiFi HotStart Uracil+ ReadyMix, 13μL eluted M-ATAC/M-CUT&Tag DNA, 1μL 10μM i5 index primer, 1μL 10μM i7 index primer) with the following PCR thermocycler program: 72 °C for 5 min; 98 °C for 30 s; 12 cycles of 98 °C for 10 s for M-ATAC or 14 cycles of 98 °C for 10 s for M-CUT&Tag, 63 °C for 30 s and 72 °C 30 s; and a final elongation at 72 °C for 1 min. DNA was purified using SPRI AMPure XP beads with a bead-to-sample ratio of 1.8:1 (no size selection) and eluted in 25 μl of H2O.

Pre-sequencing library analysis for concentration and size distribution was performed using an Agilent 2200 TapeStation with a D1000 screentape. DNA libraries were sequenced using 2x150bp paired-end reads on the NovaSeq6000 instrument.

#### RNA-seq

*Tet* KO RNA-seq was done as previously for WT and *Dnmt* KO cells^43^. Briefly, total RNA was extracted from cell pellets using the NucleoSpin RNA kit (Macherey-Nagel). Total RNA was sent to Novogene for poly(A) RNA library constructions and sequencing (see below for bioinformatic analysis of RNA-seq data) using 2x150bp paired-end reads on the NovaSeq6000 instrument.

### Bioinformatic Analysis

#### Methyl-ATAC, hMethyl-ATAC, and Methyl-CUT&Tag processing

PCR duplicate reads were filtered out before alignment using Clumpify^66^ (v38.18) with parameters ‘dedupe=t k=19 passes=6 subs=$SUBS_COUNT’, where $SUBS_COUNT corresponds to the read length multiplied by the sequencing error rate (1%). Sequencing adaptors and low-quality nucleotides were removed using Trimmomatic^67^ (v0.39) with parameters ‘HEADCROP:18 ILLUMINACLIP:adapters.fa:2:30:10 SLIDINGWINDOW:4:20 MINLEN:24’. Read quality was assessed with FastQC^68^ (v0.11.9). Processed reads were aligned to the mm10 genome using Bismark^69^ (v0.23.1) and default parameters. Reads that lost their mate pair during read trimming and/or that could not be aligned in paired-end mode were kept and re-aligned as single-end reads. Aligned reads were deduplicated with ‘deduplicate_bismark’. Methylation information of individual cytosines was extracted with ‘bismark_methylation_extractor’.

Peak calling was performed with MACS2^70^ (v2.2.7.1) and parameters ’--broad -f BAMPE -g 1.87e9 --qvalue 0.01 --nomodel --shift 0’. Genomic feature annotation was performed with ChIPseeker^71^ (v1.44.0). Promoter regions were defined as ±3 kb from the transcription start site. Regions classified as "Downstream (<=300)" and "Distal intergenic" were re-labeled as "Intergenic". Annotation of CpG islands was performed with annotatr^72^ (v1.34.0).

BigWig files of accessibility/histone mark signal were generated using deepTools^73^ (v3.5.1) bamCoverage and parameters ‘--binSize 1 --smoothLength 0 –minMappingQuality 10 --normalizeUsing CPM --blackListFileName ENCFF547MET.be --ignoreForNormalization chrX chrM chrY’. Blacklisted regions are those defined by the Kundaje lab as part of the ENCODE consortium^74^. BigWig files of 5mC levels were generated from Bismark’s CpG report by extracting the coordinates and methylation percentage of Cs covered by at least five reads. The resulting file was transformed into bigWig using bedGraphToBigWig (v4).

BigLolly (https://genome.ucsc.edu/goldenPath/help/bigLolly.html) files for 5(h)mC visualization were generated using WHAM^75^. Briefly, WHAM retrieves the methylation calls of individual (single-end or paired-end) bismark-aligned reads. This information is then transformed into a bigLolly-formatted bigBed track^76^, where each cytosine is depicted by a colored circle according to its methylation status (white - unmethylated; black - methylated). Cytosines belonging to the same read (or read pair, in the case of PE sequencing) are plotted with the same y coordinate with connecting lines, and reads are cascaded for dense viewing. Additional symbols depict the start and end of the read. To avoid overplotting, only reads with mapping quality ≥ 30 were rendered. For H3K27ac M-CUT&Tag-derived lollipops, minimum mapping quality was reduced to 10. Only a subset of lollipops is displayed in each screenshot (maximum 12 stacked reads) due to space constraints.

The UCSC Genome Browser^77^ was used for displaying NGS data, including ENCODE4 Registry of candidate Cis-Regulatory Elements (cCREs^78^). Distal enhancers (yellow): high chromatin accessibility and H3K27ac signals located farther than 2 kb from an annotated transcription start site. CA-TF (dark purple): high chromatin accessibility, low H3K4me3, H3K27ac, and CTCF signals, and transcription factor binding. CA-CTCF (blue): high chromatin accessibility and CTCF binding, but low H3K4me3 and H3K27ac signals. CA (green): high chromatin accessibility and low H3K4me3, H3K27ac, and CTCF signals.

All statistical analyses were performed on independent biological replicates. For specific applications (see below), individual replicates were merged by concatenating the corresponding .fastq files and processed as detailed above. Libraries statistics are displayed in Table S12.

#### WGBS and 5hmC-seq processing

Sequencing adaptors and low-quality nucleotides were removed using Trimmomatic^67^ with parameters ‘ILLUMINACLIP:adapters.fa:2:30:10 SLIDINGWINDOW:4:20 MINLEN:36’. Processed reads were aligned to the mm10 genome using Bismark^69^ as described above. Aligned reads were deduplicated with ‘deduplicate_bismark’. Methylation information of individual cytosines was extracted with ‘bismark_methylation_extractor’. BigWig files of 5mC levels were generated as described above.

#### RNA-seq processing

PCR duplicate reads were filtered out using Clumpify^66^ as described above. Sequencing adaptors and low-quality nucleotides were removed using Trimmomatic^67^ with parameters ‘ILLUMINACLIP:adapters.fa:2:30:10 SLIDINGWINDOW:4:20 MINLEN:24’. Reads were aligned to the mm10 genome using STAR^79^ (v2.7.9a) and parameters ’--runMode alignReads --outFilterType BySJout --outSAMtype SAM’. BigWig files were generated as described above, with the inclusion of ribosomal RNA coordinates as part of the blacklisted regions, and updated parameter ‘--minMappingQuality 255’. Read counts were obtained for each gene identifier (UCSC refGene) with htseq-count^80^ (v2.0.2) and parameters ’-s no -a 255 -m intersection-nonempty’. Differential expression was determined with DESeq2^81^ (v1.48.2) using un-normalized counts. Thresholds for significance were padj < 0.05 and |log2FoldChange| > 1. PCA was calculated from the DESeq2 output, using rlog transformation and clustering samples based on the top 500 most-variable genes.

#### Differential accessibility and differential enrichment

Differential accessibility of non-promoter ATAC peaks was performed using TCseq^82^ (v1.32.0). Identification of the five dynamic clusters was carried out with the function ’timecourseTable’, using RPKM counts normalized by effective library size. Peaks that showed no significant changes between any two time points were designated as “static” and discarded from the analysis. Temporal patterns were identified with function ‘timeclust’ and k-means clustering ranging from 3 to 8 clusters, inclusive. K = 5 was selected for further analysis, as it displayed the starkest trend differences between clusters. Differential enrichment of histone marks (H3K4me1, H3K27ac) was performed using DiffBind’s^83^ (v3.18.0) implementation of DESeq2. Peaks overlapping a promoter region were discarded from the analysis. H3K27ac peaks in closed chromatin (no overlapping ATAC peak) were filtered out likewise. Thresholds for significance were FDR < 0.05 and |log2FC| > 1.

#### Quantification of DNA (hydroxy)methylation

5(h)mC was calculated using merged replicates to achieve higher read coverage. Only CpGs covered by at least 5 reads were considered. For boxplots representing the levels of 5mC or 5hmC from M-ATAC versus hM-ATAC samples (Figure 5A), one set of peaks was used per condition (two merged replicates for WT and one replicate for KO ExEpiLC controls). Average 5(h)mC reported per locus was calculated from the 5(h)mC of each individual CpG, weighted by the read coverage of the nucleotide. Average 5(h)mC at enhancers (see enhancer classification below) was quantified over the entire enhancer coordinate, irrespective of the ATAC/H3K27ac peak coordinate. For global 5mC levels (Figure S2G), the non-weighted average was reported. Due to incomplete enzymatic conversion of the H3K27ac Methyl-CUT&Tag samples (mean conversion rate = 86.8%), reads that exhibited non-CpG methylation (which is known to be depleted in mammalian genomes) were ignored during 5mC quantification. Differentially methylated regions were identified with MethyLasso^84^ (v1.0.0) using default parameters. The input files were obtained with Bismark’s^69^ function ‘coverage2cytosine’ and parameter ’--merge_CpG’, followed by removal of cytosines located in non-canonical chromosomes (chrUn_* and chr*_random).

#### Enhancer classification

For every combination of cell type (mESCs, EpiLCs, ExEpiLCs) and chromatin feature (H3K4me1, H3K27ac, ATAC), a "master" peak set was generated from WT data. A peak was part of the master set if: 1) it was called in the dataset generated by merging the replicates; and 2) overlapped with a peak identified in at least one replicate. These two criteria were selected to balance the presence of false negatives, as more stringent filters tended to misclassify active and primed enhancers. Each peak of the H3K4me1 master set was annotated as: 1) active, if it overlapped a master ATAC peak and a master H3K27ac peak; 2) primed, if it overlapped a master ATAC peak but not a master H3K27ac peak; or 3) inaccessible, if it did not overlap neither a master ATAC peak nor a master H3K27ac peak. The three H3K4me1 master peak sets (one per time point) were combined using BEDTools^85^ (v2.31.1) ‘merge’ function to generate a "base" set of enhancer coordinates. Each coordinate was subsequently annotated based on the cell type-specific H3K4me1 classification. Enhancers that overlapped a promoter region were discarded. Association of enhancers to nearby genes was performed using GREAT^50,86^ (v4.0.4) with default parameters. Enrichment analysis of mapped genes was performed with clusterProfiler^87^ (v4.16.0) and visualized with enrichplot^88^ (v1.28.4). Gene ontologies depicted belong to "Biological Process."

#### Motif analysis

TF motif identification was performed with TFmotifView^89^ using the 2026 JASPAR database^90^. Non-mouse TFs were associated to their *Mus musculus* homolog using biomaRt^91,92^ (v2.64.0). Motifs corresponding to non-expressed TFs (< 1 normalized transcript read) were discarded from the analysis. A motif was determined to be significantly enriched if its hypergeometric p-value was below the probability of occurrence of a random hexamer (0.25^6). TF sensitivity to 5mC was extracted from MethylSelex^5^. TFs identified both as "MethylPlus" and "MethylMinus" are referred to as "Dual" in this publication. Motifs were annotated as CpG-containing if their degenerate consensus sequence contained: 1) a CpG dinucleotide; 2) a C followed by a degenerated G [RSKVDBN]; 3) a degenerated C [MSYVHBN] followed by a G; or 4) were assessed by MethylSelex^5^ despite not having a major CG in the degenerate consensus motif. For motif identification at the five dynamic ATAC clusters, static peaks were used as background. For motif identification at 5hmC-enriched regions, the top 10% of hM-ATAC peaks (5hmC % ≥ 17.6%) were compared against the bottom 10% hM-ATAC peaks (5hmC % ≤ 0.14%).

## Supplemental tables

Table S1: Number of differentially accessible M-ATAC peaks per cluster (related to Figure 2B).

Table S2: Top 20 ontologies of the genes associated with the 5 clusters of dynamic peaks (related to Figure 2B).

Table S3: Statistical difference in 5mC (related to Figure 2C,E and Figure S2D).

Table S4: Number of differential peaks per cluster (related to Figure 3, Figure S3, and Figure S4).

Table S5: p-values of 5mC changes at differential H3K27ac (related to Figure 4E and Figure S6B).

Table S6: p-values of 5hmC in DMRs (related to Figure 5B) and enhancer classes (related to Figure 5D).

Table S7: p-values of RNA changes linked to differential H3K27ac (related to Figure 6D and Figure S9A,B).

Table S8: TF motifs found at the 5 clusters of dynamic peaks (related to Figure S10A).

Table S9: TF motifs found at differentially accessible peaks between WT and *Dnmt* KO cells (related to Figure 7B and Figure S10B).

Table S10: Lists of motifs found at the 5hmC enriched hM-ATAC peaks (related to Figure 7C and Figure S10C).

Table S11: List of the sequence of modified oligonucleotides and primers. Table S12: Libraries statistics.

---

**Figure S1.**
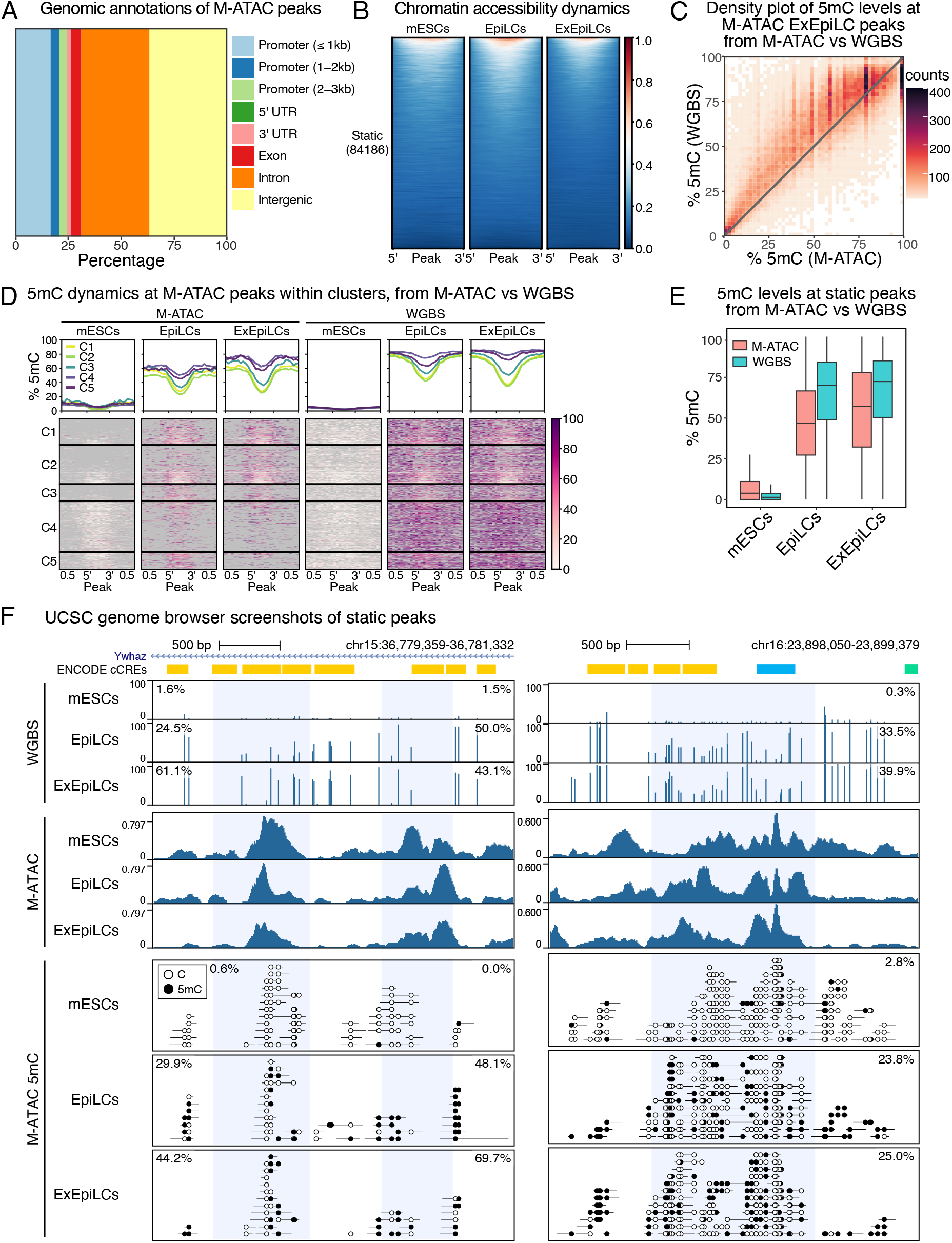
(related to. Figure 1**): A.** Genomic annotations of M-ATAC peaks (all WT samples merged). **B.** Heatmaps showing the M-ATAC signal within static non-promoter peaks in WT cells during mESC to EpiLC to ExEpiLC differentiation (84186 peaks). **C.** Density plot of average 5mC from M-ATAC non-promoter peaks compared with WGBS for ExEpiLCs (84823 peaks, Pearson correlation = 0.81). **D.** Heatmaps showing WT 5mC signal from M-ATAC and WGBS during mESC to EpiLC to ExEpiLC differentiation within the 5 clusters shown in Figure 1C. **E.** Boxplots of 5mC percentages from M-ATAC (red) and WGBS (blue) in mESCs, EpiLCs, and ExEpiLCs, for the static peaks presented in panel B. **F.** Representative UCSC genome browser screenshots of M-ATAC and WGBS at two putative static enhancer regions. Blue shading indicates the putative enhancer region defined by our analysis. A description of different ENCODE cCREs can be found in Methods. Methylation percentages of the highlighted regions for WGBS and M-ATAC are indicated.

**Figure S2.**
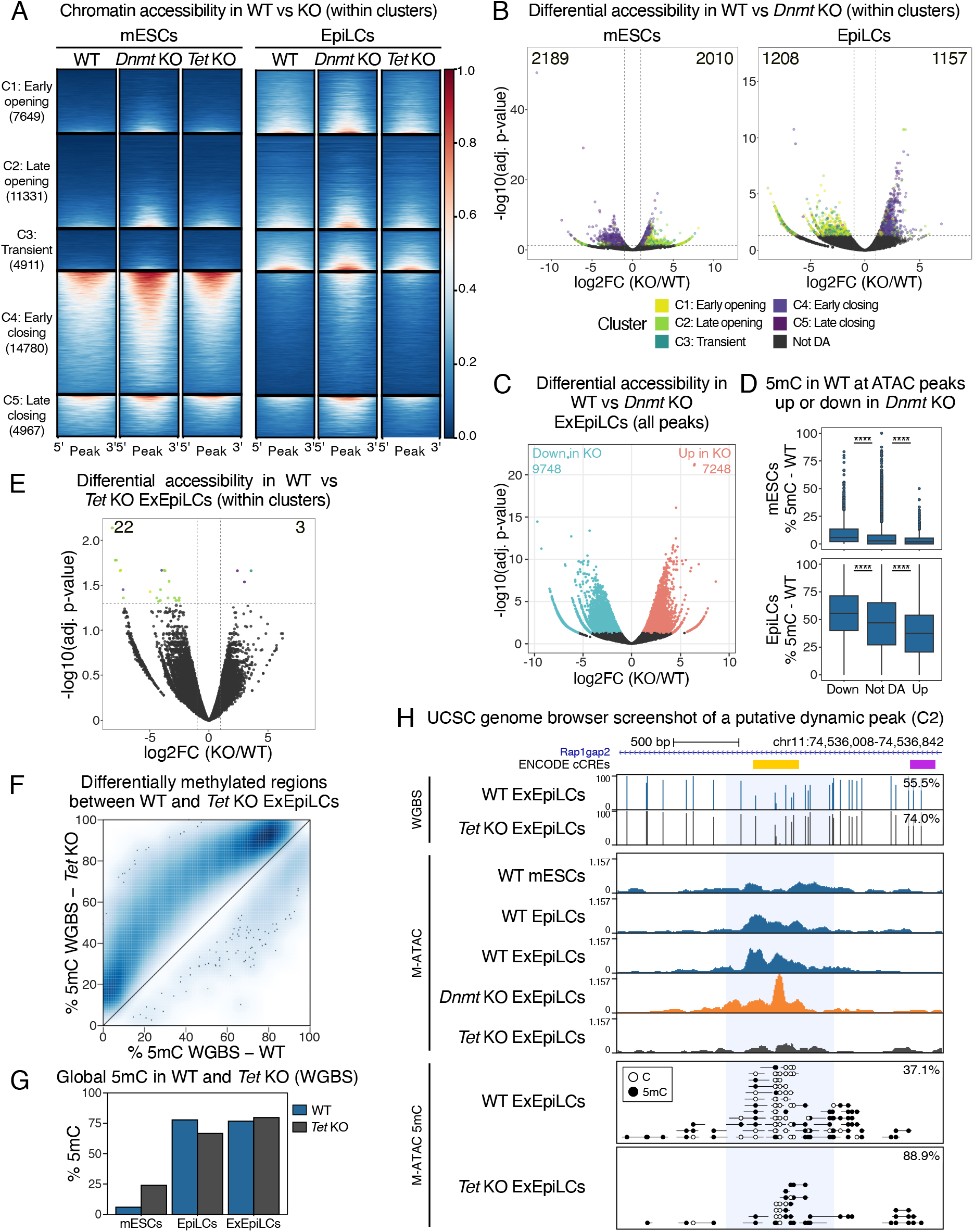
(related to. Figure 2**): A.** Heatmaps showing the M-ATAC signal in WT, *Dnmt* KO, and *Tet* KO mESCs (left) and EpiLCs (right), within the 5 clusters defined in Figure 1C. **B.** Volcano plots of differential chromatin accessibility of peaks within the 5 clusters between WT and *Dnmt* KO mESCs (left) and EpiLCs (right) (log2-fold change > 1 or < -1; adjusted p-value < 0.05). Peaks are colored based on their cluster affiliation. The number of less accessible (down) and more accessible (up) peaks is indicated. **C.** Volcano plot of differential chromatin accessibility of all M-ATAC non-promoter peaks (log2-fold change > 1 or < -1; adjusted p-value < 0.05). **D.** Boxplots of 5mC levels in WT mESCs (top) and EpiLCs (bottom), comparing the peaks that are down, up, or not differentially accessible (not DA) in *Dnmt* KO compared to WT. **E.** Volcano plot of differential chromatin accessibility between WT and *Tet* KO ExEpiLCs at peaks within the 5 clusters. **F.** Differential methylated regions between WT and *Tet* KO ExEpiLCs from WGBS. **G.** Global 5mC levels in WT (blue) versus *Tet* KO (grey) mESCs, EpiLCs, and ExEpiLCs from WGBS. **H.** Representative UCSC genome browser screenshot of M-ATAC and WGBS at a putative enhancer region associated with the late opening cluster (C2). White lollipops represent unmethylated cytosines and black lollipops represent methylated cytosines. Blue shading indicates the putative enhancer region defined by our analysis. A description of different ENCODE cCREs can be found in Methods. Methylation percentages of the highlighted region are indicated. *p < 0.05; **p < 0.01; ***p < 0.001; ****p < 0.0001; two-sided Wilcoxon test with Bonferroni correction.

**Figure S3.**
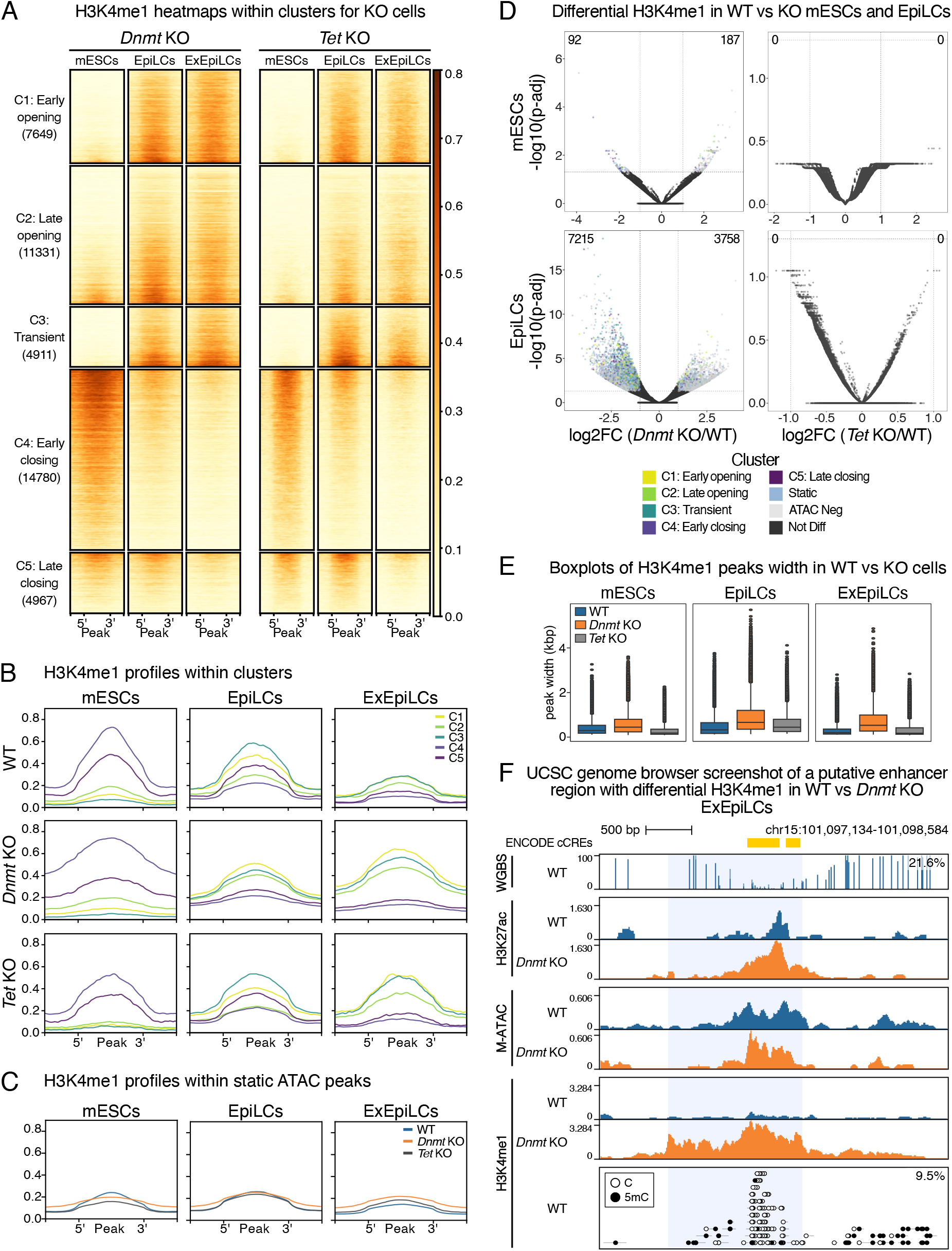
(related to. Figure 3**): A.** Heatmaps showing the H3K4me1 M-CUT&Tag signal for *Dnmt* KO (left) and *Tet* KO (right) (as comparison to WT in Figure 3B) within the 5 clusters defined in Figure 1C. **B.** Average profiles of H3K4me1 corresponding to the heatmaps shown in Figure 3B for WT and Figure S3A for *Dnmt* KO and *Tet* KO. **C.** Average profiles of H3K4me1 at static peaks for WT, *Dnmt* KO, and *Tet* KO cells. **D.** Volcano plots showing differential enrichment of H3K4me1 in WT vs *Dnmt* KO (left) or *Tet* KO (right) cells (top: mESCs, bottom: EpiLCs) (log2-fold change > 1 or < -1; adjusted p-value < 0.05). Peaks are colored based on their overlap with ATAC-accessible regions (dynamic or static clusters) or lack thereof (ATAC Neg). **E.** Boxplots of H3K4me1 peaks width in WT, *Dnmt* KO, and *Tet* KO in mESCs (left), EpiLCs (middle), and ExEpiLCs (right). **F.** Representative UCSC genome browser screenshot of putative enhancer regions (ENCODE cCREs) with changing H3K4me1 in *Dnmt* KO compared to WT ExEpiLCs. White lollipops represent unmethylated cytosines and black lollipops represent methylated cytosines. Blue shading indicates the putative enhancer region defined by our analysis. Methylation percentages of the highlighted region are indicated.

**Figure S4.**
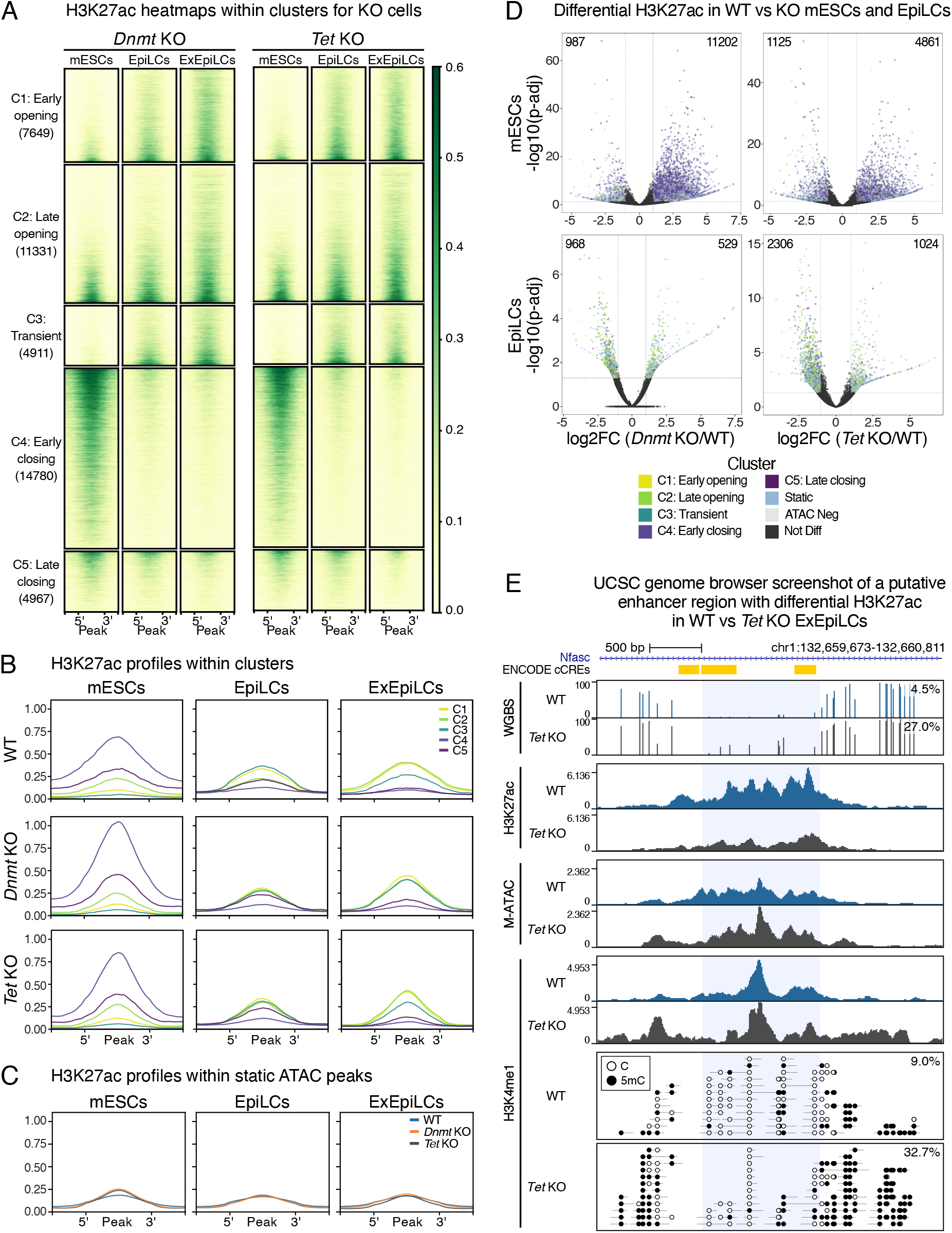
(related to. Figure 3**): A.** Heatmaps showing the H3K27ac M-CUT&Tag signal for *Dnmt* KO (left) and *Tet* KO (right) (as comparison to WT in Figure 3B) within the 5 clusters defined in Figure 1C. **B.** Average profiles of H3K27ac corresponding to the heatmaps shown in Figure 3B for WT and Figure S4A for *Dnmt* KO and *Tet* KO. **C.** Average profiles of H3K27ac at static peaks for WT, *Dnmt* KO, and *Tet* KO. **D.** Volcano plots showing differential enrichment of H3K27ac in WT vs *Dnmt* KO (left) or *Tet* KO (right) cells (top: mESCs, bottom: EpiLCs) (log2-fold change > 1 or < -1; adjusted p-value < 0.05). Peaks are colored based on their overlap with ATAC-accessible regions (dynamic or static clusters) or lack thereof (ATAC Neg). **E.** Representative UCSC genome browser screenshot of putative enhancer regions (ENCODE cCREs) with changing H3K27ac in *Tet* KO compared to WT ExEpiLCs. White lollipops represent unmethylated cytosines and black lollipops represent methylated cytosines. Blue shading indicates the putative enhancer region defined by our analysis. Methylation percentages of the highlighted region are indicated.

**Figure S5.**
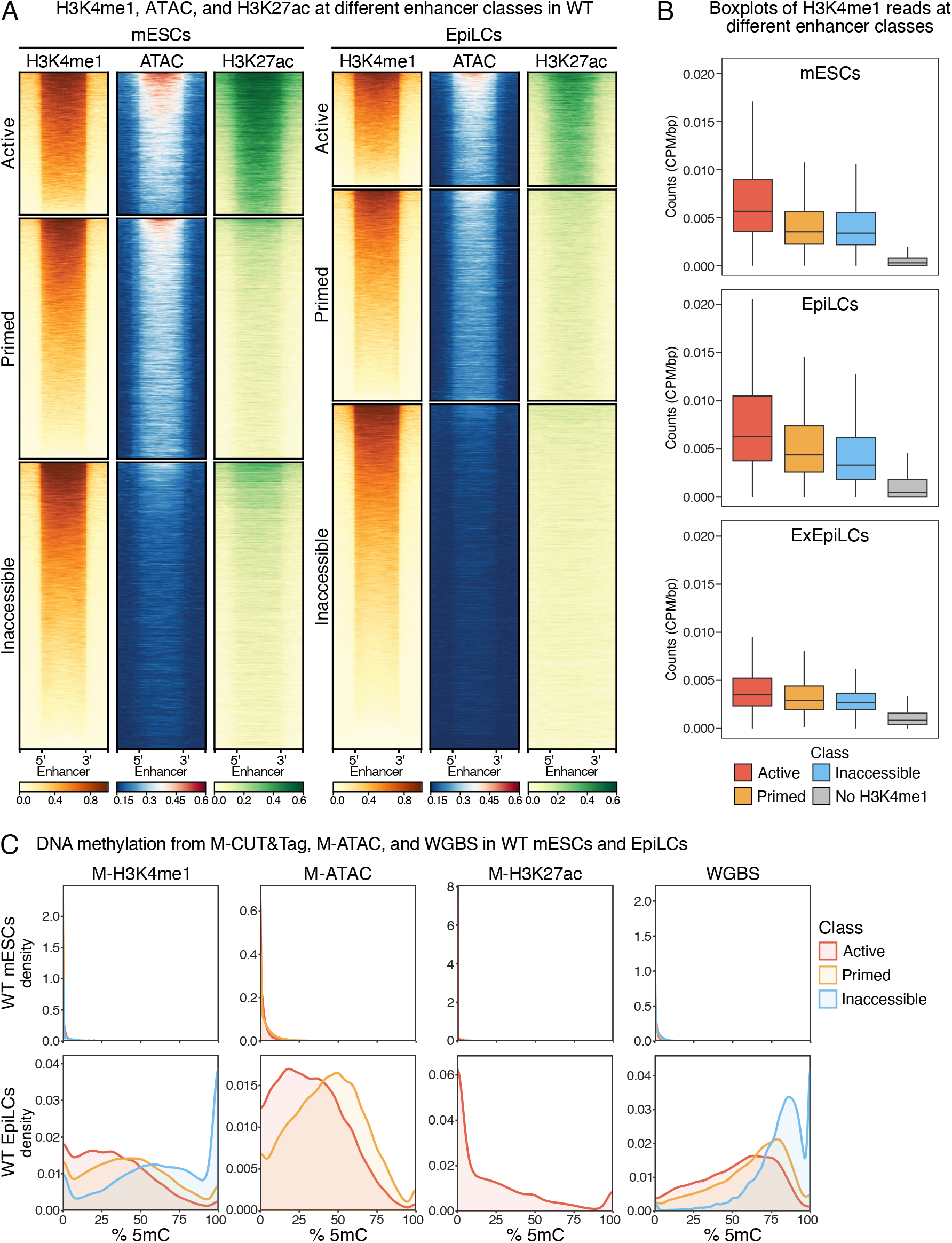
(related to. Figure 4**): A.** Heatmaps showing the H3K4me1, M-ATAC, and H3K27ac signal in WT mESCs (left panels) and EpiLCs (right panels) for enhancers classified as: active (positive for all three features), primed (positive for H3K4me1 and ATAC and negative for H3K27ac) and inaccessible enhancers (positive for H3K4me1 and negative for ATAC and H3K27ac). **B.** Boxplots of WT H3K4me1 signal at different enhancer classes. H3K4me1 signal was normalized by library size (CPM) and enhancer length. Enhancers identified as such in at least another time point, but without an associated H3K4me1 peak in the corresponding day, are depicted in grey. **C.** Density plots of 5mC levels at different classes of enhancers, quantified from M-ATAC, H3K4me1 and H3K27ac M-CUT&Tag, and WGBS datasets in WT mESCs (top) and EpiLCs (bottom).

**Figure S6.**
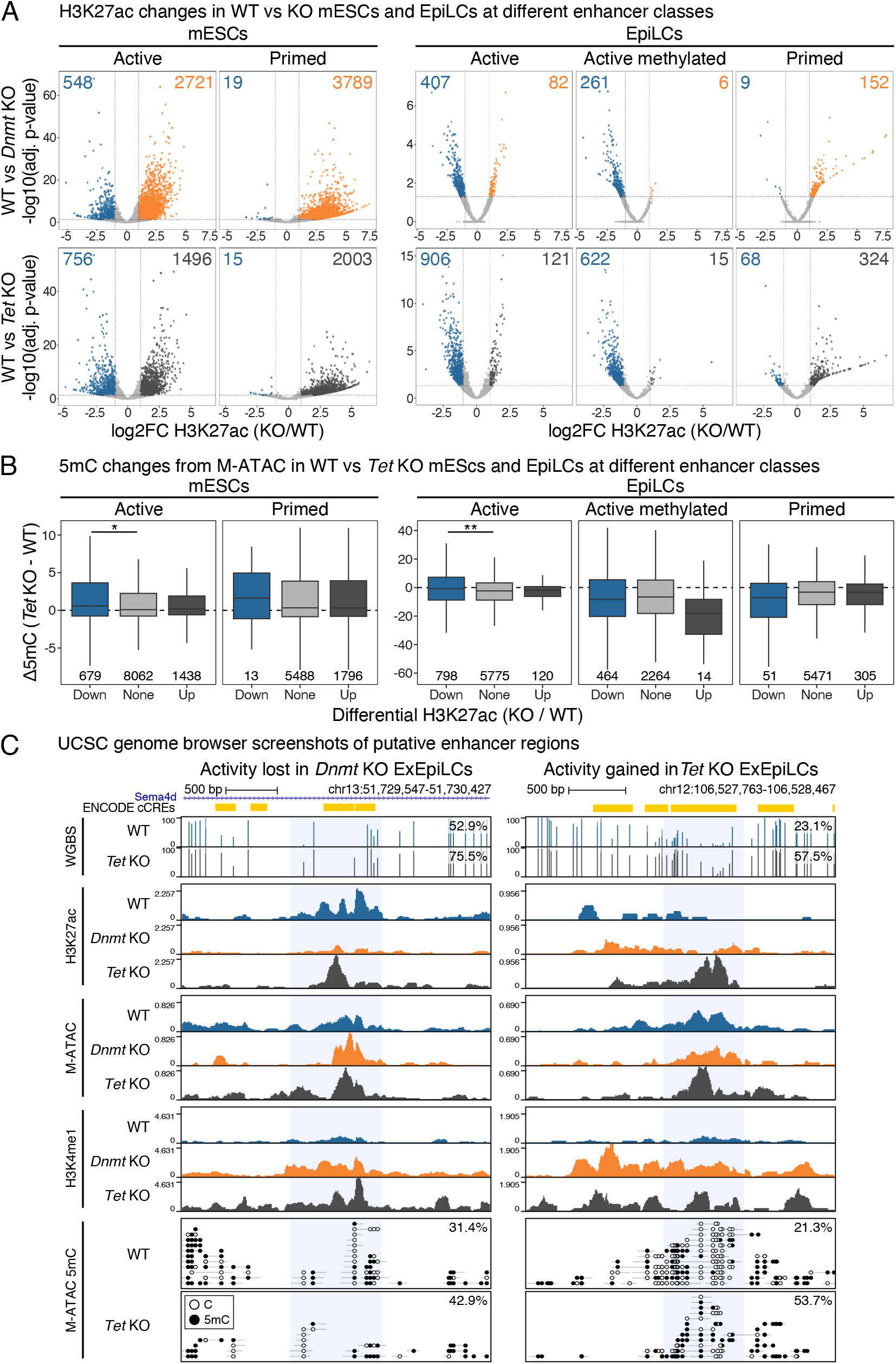
(related to. Figure 4**): A.** Volcano plots showing differential enrichment of H3K27ac in WT vs *Dnmt* KO (top) or *Tet* KO (bottom) at the different classes of enhancers (active, active methylated, and primed) (log2-fold change > 1 or < -1; adjusted p-value < 0.05). mESCs are shown in left and EpiLCs are shown in right panels. **B.** 5mC changes quantified from M-ATAC datasets at enhancers associated with gain or loss of H3K27ac in WT versus *Tet* KO cells, separated by enhancer class. mESCs are shown in left and EpiLCs are shown in right panels. The number of enhancers per box is indicated below. **C.** Representative UCSC genome browser screenshots of putative active methylated (left) and primed (right) enhancer regions (ENCODE cCREs). White lollipops represent unmethylated cytosines and black lollipops represent methylated cytosines. Blue shading indicates the putative enhancer region defined by our analysis. Methylation percentages of the highlighted regions are indicated. *p < 0.05; **p < 0.01; ***p < 0.001; ****p < 0.0001; two-sided Wilcoxon test with Bonferroni correction.

**Figure S7.**
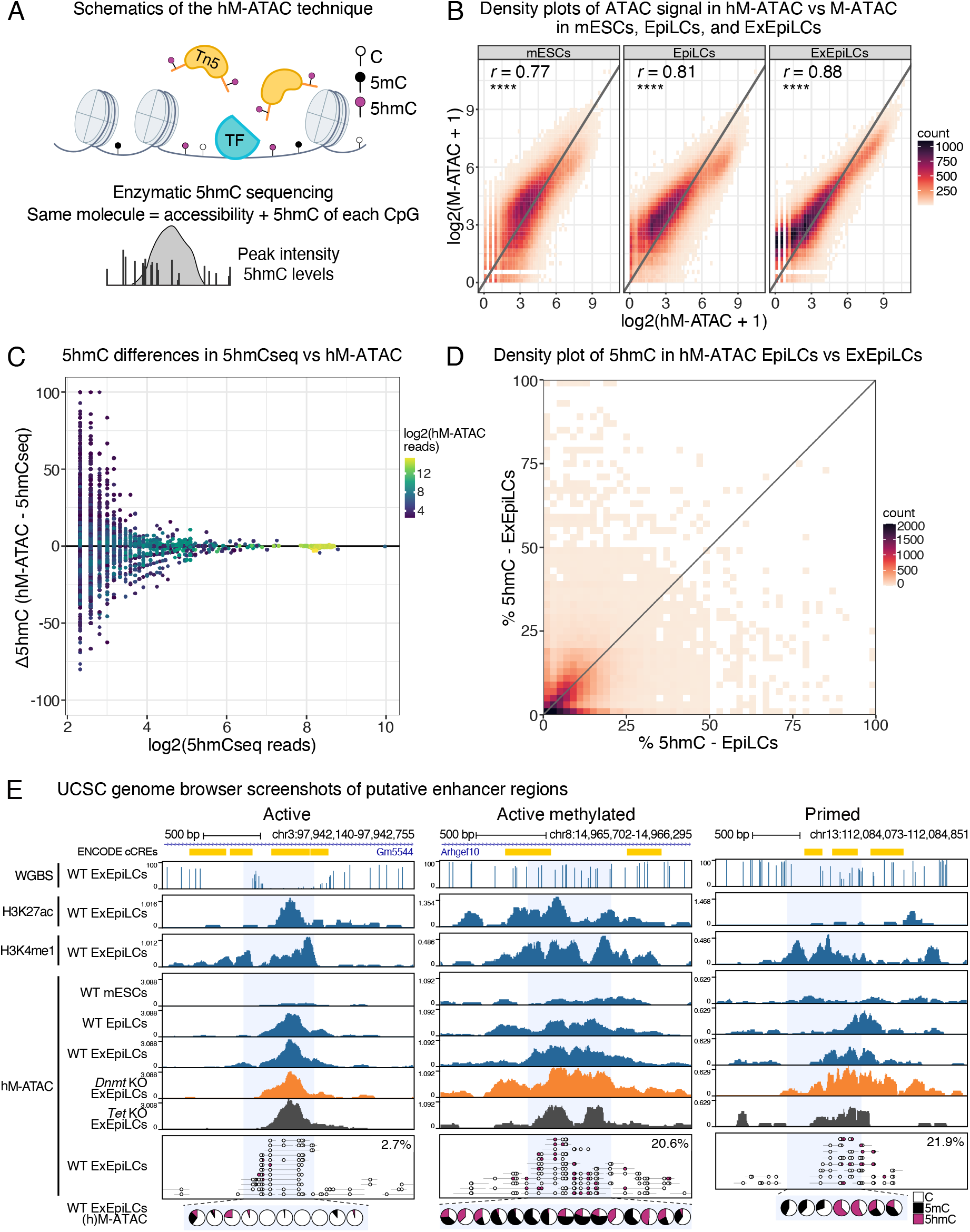
(related to. Figure 5**): A.** Schematic overview of the hM-ATAC technique. **B.** Density plots of WT ATAC signal in hM-ATAC vs M-ATAC in mESCs, EpiLCs, and ExEpiLCs (r = Pearson correlation). **C.** Accuracy of 5hmC quantification versus sequencing depth. Delta of 5hmC levels between hM-ATAC and 5hmCseq (Y axis) plotted against 5hmCseq coverage (X axis) for individual CpGs in WT ExEpiLCs. Color represents coverage in hM-ATAC dataset. **D.** Density plot of average 5hmC levels in hM-ATAC EpiLCs vs ExEpiLCs (63514 peaks, Pearson correlation = 0.29). **E.** UCSC genome browser screenshots of putative enhancer regions (ENCODE cCREs). White and magenta lollipops represent unmethylated and hydroxymethylated cytosines, respectively. Blue shading indicates the putative enhancer region defined by our analysis. Hydroxymethylation percentages of the highlighted regions are indicated. The pie charts represent the proportion of C, 5mC, and 5hmC at each CpG in the shaded window, inferred from M- and hM-ATAC. *p < 0.05; **p < 0.01; ***p < 0.001; ****p < 0.0001.

**Figure S8.**
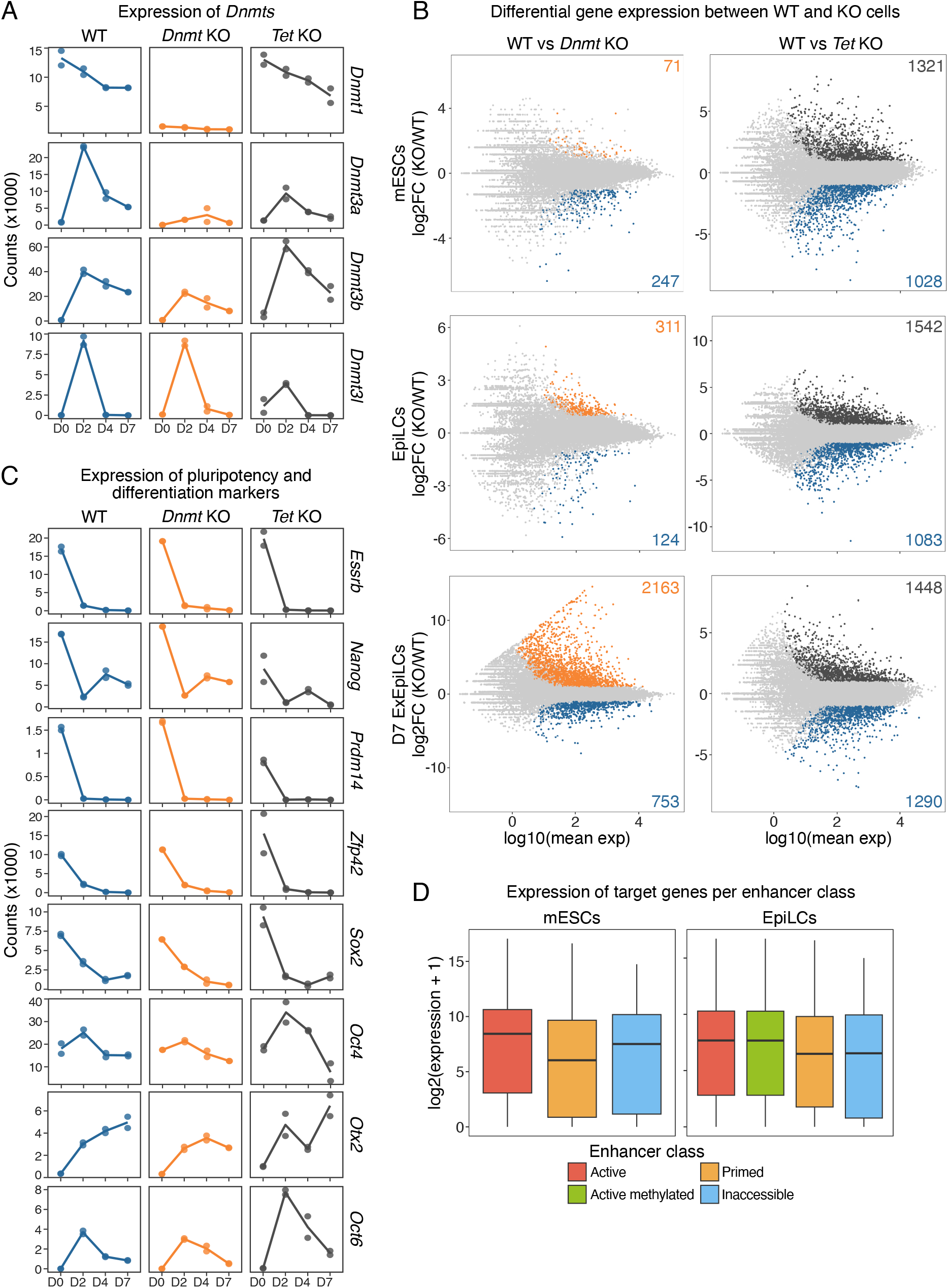
(related to. Figure 6**): A.** Normalized read counts of *Dnmt* genes from RNA-seq in WT and KO mESCs (D0), EpiLCs (D2), ExEpiLCs (D4), and ExEpiLCs after 7 days of differentiation (D7). Solid line represents the mean of biological duplicates. **B.** MA plots of differential gene expression in WT vs *Dnmt* KO (left) and WT vs *Tet* KO (right) cells (mESCs, EpiLCs, and D7 ExEpiLCs). The number of up-regulated and down-regulated genes is indicated. **C.** Normalized read counts of naive pluripotency (*Essrb, Nanog, Prdm14, Zfp42*), general pluripotency (*Sox2, Oct4*) and formative/primed pluripotency (*Otx2, Oct6*) markers from RNA-seq in WT and KO cells. **D.** Normalized read counts of genes associated with each enhancer class (active, active methylated, primed, and inaccessible) in WT mESCs (left) and EpiLCs (right).

**Figure S9.**
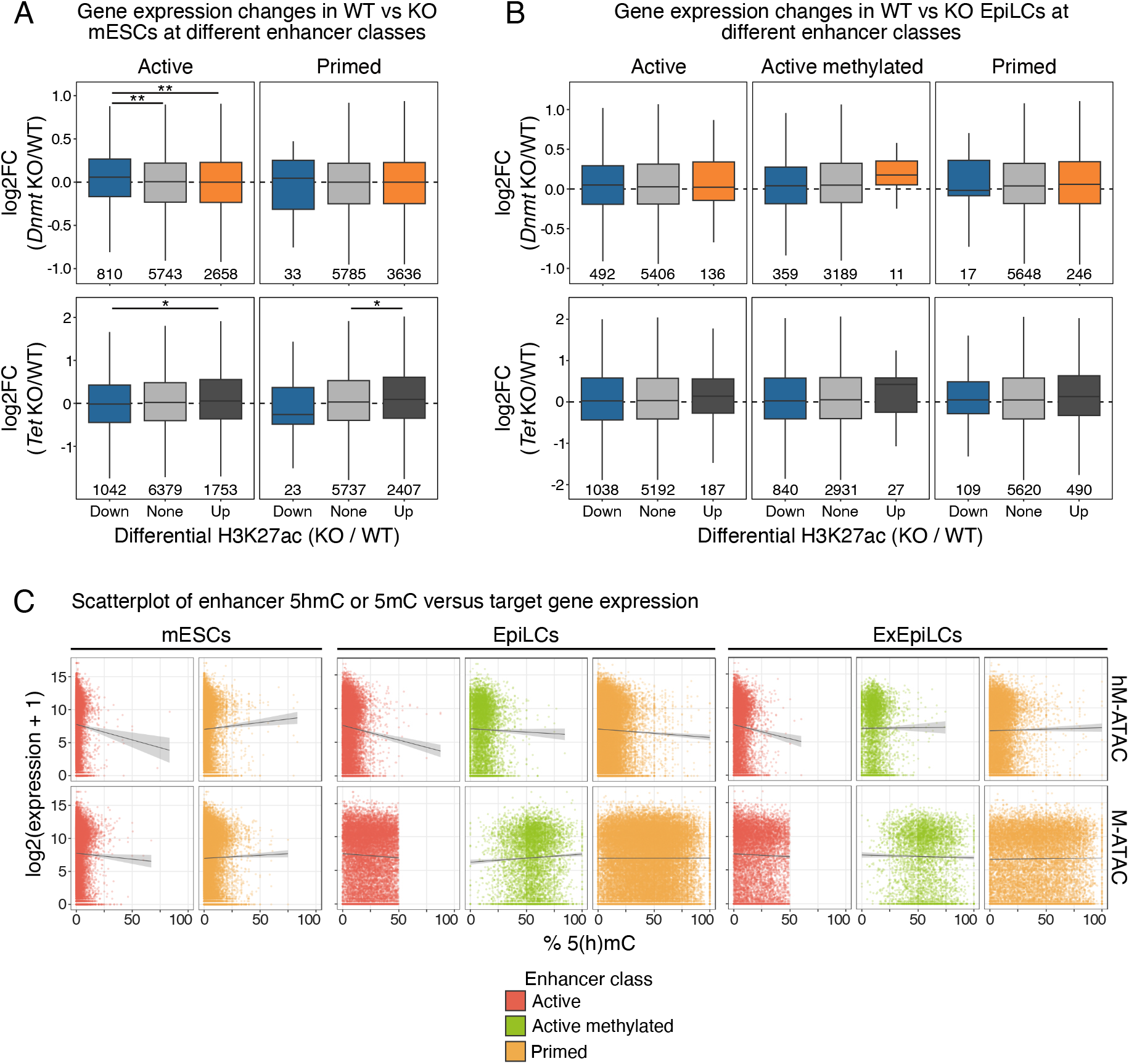
(related to. Figure 6**): A,B.** RNA expression differences in genes associated with enhancers displaying gain/loss of H3K27ac in WT vs KO mESCs **(A)** or EpiLCs **(B)**, separated by enhancer class (active, active methylated, and primed). The number of genes per box is indicated below. *p < 0.05; **p < 0.01; ***p < 0.001; ****p < 0.0001; two-sided Wilcoxon test with Bonferroni correction. **C.** Enhancer DNA (hydroxy)methylation (X axis; 5hmC top, 5mC bottom) versus normalized read counts of their associated genes (Y axis), separated by enhancer class and cell type. Linear fit with 95% confidence intervals is depicted in grey.

**Figure S10.**
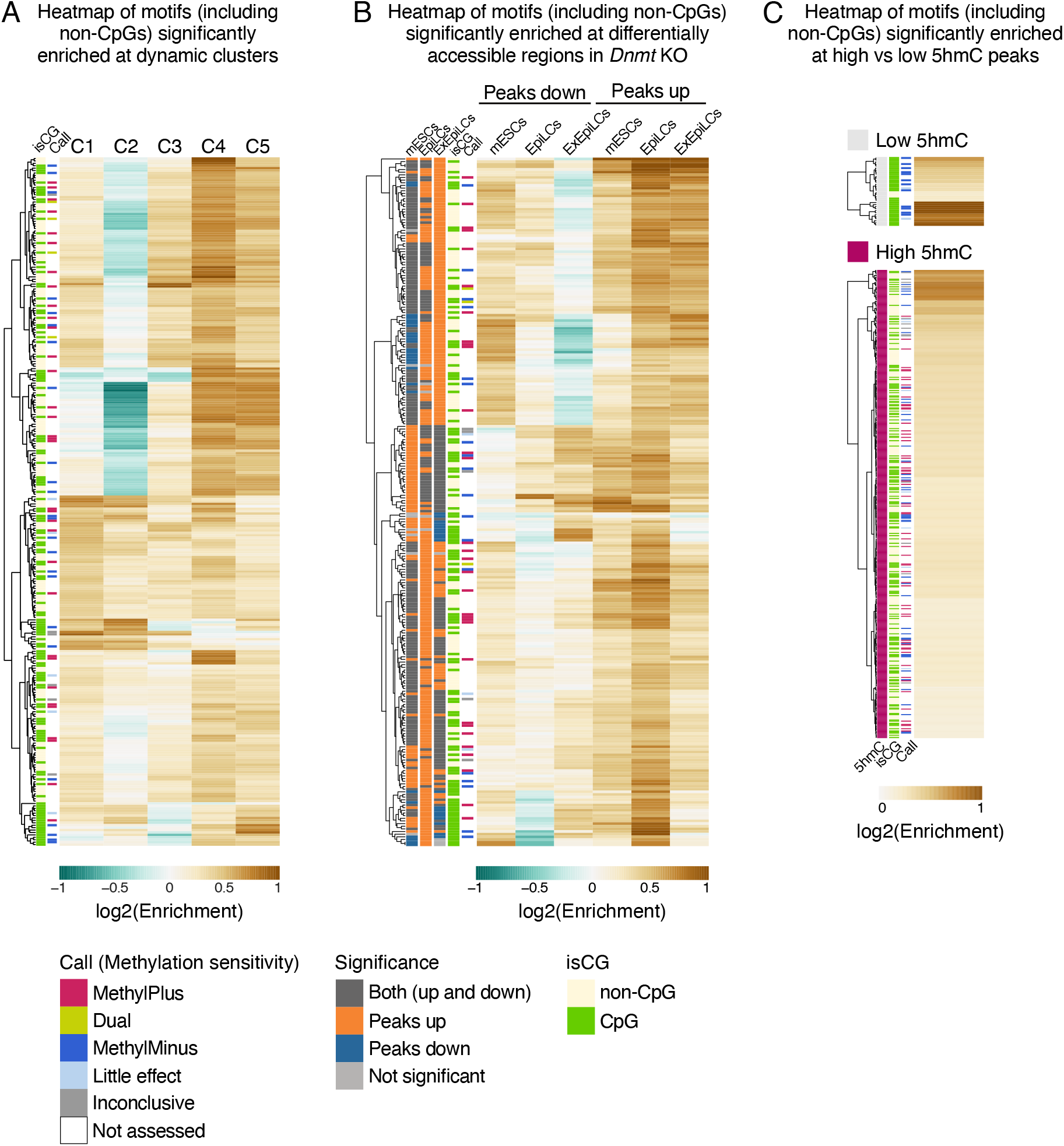
(related to. Figure 7**): A.** Heatmap of motifs significantly enriched at the five clusters of dynamic enhancers (related to Figure 1C). **B.** Same as Figure 7B, but including motifs without characterized methylation sensitivity or without a CpG in the consensus sequence. **C.** Same as Figure 7C, but including motifs without characterized methylation sensitivity or without a CpG in the consensus sequence.

